# Crosstalk between microRNA expression and DNA methylation drive the hormone-dependent phenotype of breast cancer

**DOI:** 10.1101/2020.04.12.038182

**Authors:** Miriam Ragle Aure, Thomas Fleischer, Sunniva Bjørklund, Jørgen Ankill, Jaime A. Castro-Mondragon, OSBREAC, Anne-Lise Børresen-Dale, Kristine K. Sahlberg, Anthony Mathelier, Xavier Tekpli, Vessela N. Kristensen

## Abstract

**Background:** Abnormal DNA methylation is observed as an early event in breast carcinogenesis. However, how such alterations arise is still poorly understood. microRNAs (miRNAs) regulate gene expression at the post-transcriptional level and have been shown to play key roles in various biological processes. Here, we integrate miRNA expression and DNA methylation at CpGs to study how miRNAs may affect the breast cancer methylome and how DNA methylation may regulate miRNA expression.

**Results:** miRNA expression and DNA methylation data from two breast cancer cohorts were subjected to genome-wide correlation analysis. Clustering of the miRNA expression-DNA methylation association pairs significant in both cohorts identified distinct clusters of miRNAs and CpGs. These clusters recapitulated important biological processes associated with breast cancer pathogenesis. Notably, two major clusters were related to immune or fibroblast infiltration, hence identifying miRNAs associated with cells of the tumor microenvironment, while another large cluster was related to estrogen receptor (ER) signaling. Studying the chromatin landscape surrounding the CpGs associated with the estrogen-signaling cluster, we found that miRNAs from this cluster are likely to be regulated through DNA methylation of enhancers bound by FOXA1, GATA2 and ER-alpha. Further, at the hub of the estrogen-cluster, we identified hsa-miR-29c-5p as negatively correlated with the mRNA and protein expression of the DNA methyltransferase DNMT3A, a key enzyme regulating DNA methylation. We found deregulation of hsa-miR-29c-5p already in pre-invasive breast lesions and postulate that hsa-miR-29c-5p may trigger early event abnormal DNA methylation in ER positive breast cancer.

**Conclusions:** We describe how miRNA expression and DNA methylation interact and associate with distinct breast cancer phenotypes.

## Background

Breast cancers are highly heterogeneous at the clinical and molecular level. Alterations of methylation at CpGs are found already in breast pre-invasive lesions [1] and are thought to shape the methylation patterns found in the different clinical and molecular breast cancer subtypes [2, 3]. The epigenome contributes to the cancer cells’ phenotype by regulating gene expression and the accessibility of regulatory regions. Previous studies have identified aberrant DNA methylation at gene promoters in breast cancer associated with clinically relevant subgroups. However, little attention has been given to DNA methylation at other non-coding regions. We recently showed that DNA methylation at enhancers identifies distinct breast cancer lineages [2]. The epigenome and the chromatin landscape are important features to explain breast cancer development and also progression, as recently demonstrated for endocrine resistance in breast cancer [4]. It is therefore essential to understand the crosstalk between the genome and the epigenome and its role in defining tumor phenotypes. Key enzymes, such as DNA methyltransferases (DNMTs) and Ten-eleven translocation enzymes (TETs) regulate the DNA methylation machinery and alterations of their expressions have been described in cancers with serious consequences in terms of cancer cell phenotype [5, 6]. However, how such enzymes may be early deregulated during carcinogenesis is still unclear.

MicroRNAs (miRNAs) are a type of small (∼22 nucleotides) non-coding RNAs regulating protein expression through targeting of messenger RNA (mRNA) for degradation or by inducing translational repression [7]. miRNAs play crucial roles in the regulation of cancer-associated processes such as proliferation, apoptosis, and differentiation, and are known to elicit context- and cell-type specific expression [8, 9]. In breast cancer, expression of miRNAs has been associated with clinical and molecular subtypes [10-12], progression [13-15], prognosis [16, 17] and expression of oncogenes [18]. Importantly, miRNAs have been shown to regulate the expression of epigenetic regulators such as DNMTs and TETs [19, 20]. We have previously shown how concerted alterations in copy number or promoter methylation affect miRNA expression *in cis*, resulting in up-regulation of oncogenic miRNAs and down-regulation of tumor-suppressor miRNAs [21]. However, how DNA methylation at distal regulatory regions is associated with miRNA expression in breast cancer remains poorly understood.

The aim of this study was to elucidate how miRNA expression associates with genome-wide DNA methylation patterns in breast cancer *in cis* and *in trans*. Specifically, we studied the interplay between miRNA expression and DNA methylation and how it may impact breast cancer subtypes. To this end, we integrated whole-genome miRNA expression with CpG DNA methylation and performed a genome-wide correlation analysis identifying <u>mi</u>RNA-<u>m</u>ethylation <u>Q</u>uantitative <u>T</u>rait <u>L</u>oci (mimQTLs). We combined and integrated mimQTLs with mRNA/protein expression, clinical information, genome segmentation, Assay for Transposase-Accessible Chromatin using sequencing (ATAC-seq) data, transcription factor (TF) binding, and Chromatin Interaction Analysis by Paired-End Tag Sequencing (ChIA-PET) RNA polymerase II (Pol2) data to elucidate the miRNA-methylation crosstalk in breast cancer.

## Results

### Identification of miRNA-methylation Quantitative Trait Loci (mimQTLs)

To identify robust associations between the expression of miRNAs and DNA methylation at CpG sites, we correlated genome-wide miRNA expression and DNA methylation in two independent breast cancer cohorts: Oslo2 (n=297) and The Cancer Genome Atlas (TCGA) Breast Invasive Carcinoma cohort (n=439; see Additional file 1 for workflow outline). Only miRNAs and CpGs found in both cohorts were considered for the estimation of the Spearman correlation between the expression of 346 miRNAs and methylation of 142 804 CpGs, resulting in 140 443 (0.28%) and 1 351 887 (2.74%) significant miRNA-CpG associations (Bonferroni-corrected p-value<0.05) in the Oslo2 and TCGA cohorts, respectively (see Methods). With a greater sample size for TCGA, a larger number of significant associations were observed, as expected (Additional file 2). We identified 89 118 significant correlations with the same sign in both cohorts, being the first evidence of robust associations between miRNA expression and DNA methylation (Additional file 3). These correlations involved 119 unique miRNAs and 26 746 unique CpGs (Additional file 4a, b). A significant correlation between the expression of a miRNA and methylation at a CpG is hereafter referred to as a *<u>mi</u>RNA-<u>m</u>ethylation <u>Q</u>uantitative <u>T</u>rait <u>L</u>oci* (mimQTL).

### Identification of mimQTL clusters

To identify mimQTLs sharing similar features of biological relevance in breast cancer, we performed unsupervised hierarchical clustering of the Spearman correlation p-values binarized as −1 (negative) and +1 (positive), which led to the identification of three miRNA clusters (x-axis) and two CpG clusters (y-axis) (Figure 1a). miRNA clusters A, B, and C consisted of 23, 59, and 37 miRNAs, respectively. CpG clusters 1 and 2 contained 14 040 and 12 706 CpGs, respectively. For each miRNA cluster, the number of associated mimQTL pairs were 66 202 (74% in cluster A), 9252 (11% in B) and 13 664 (15% in C), respectively. We observed more negative correlations (64%) than positive ones (Additional file 5). Of the CpGs associated with miRNAs in cluster C, 60% were also associated with miRNAs in cluster A. However, the sign of the correlation was inverted as shown by the opposite blue or red colors on the heatmap (Figure 1a). In contrast, most of the CpGs associated with miRNAs in cluster B were unique to this cluster. The number of CpG associations *per* miRNA varied from one up to 14 469 (hsa-miR-155-5p) with a median of 79 CpG associations (Additional file 4a and Additional file 6a). For the CpGs, the number of associations to miRNAs ranged from one up to 30 with a median of three; only a small subset (1.2%) of almost exclusively cluster 1 CpGs had more than 10 miRNA associations (Additional file 4b and Additional file 6b). As expected, a high degree of co-expression and co-methylation was observed by the members of a given cluster (Additional file 7). These initial analyses led us to identify for the first time global and robust correlations between miRNA expression and CpG methylation across two breast cancer cohorts.

**Figure 1.**
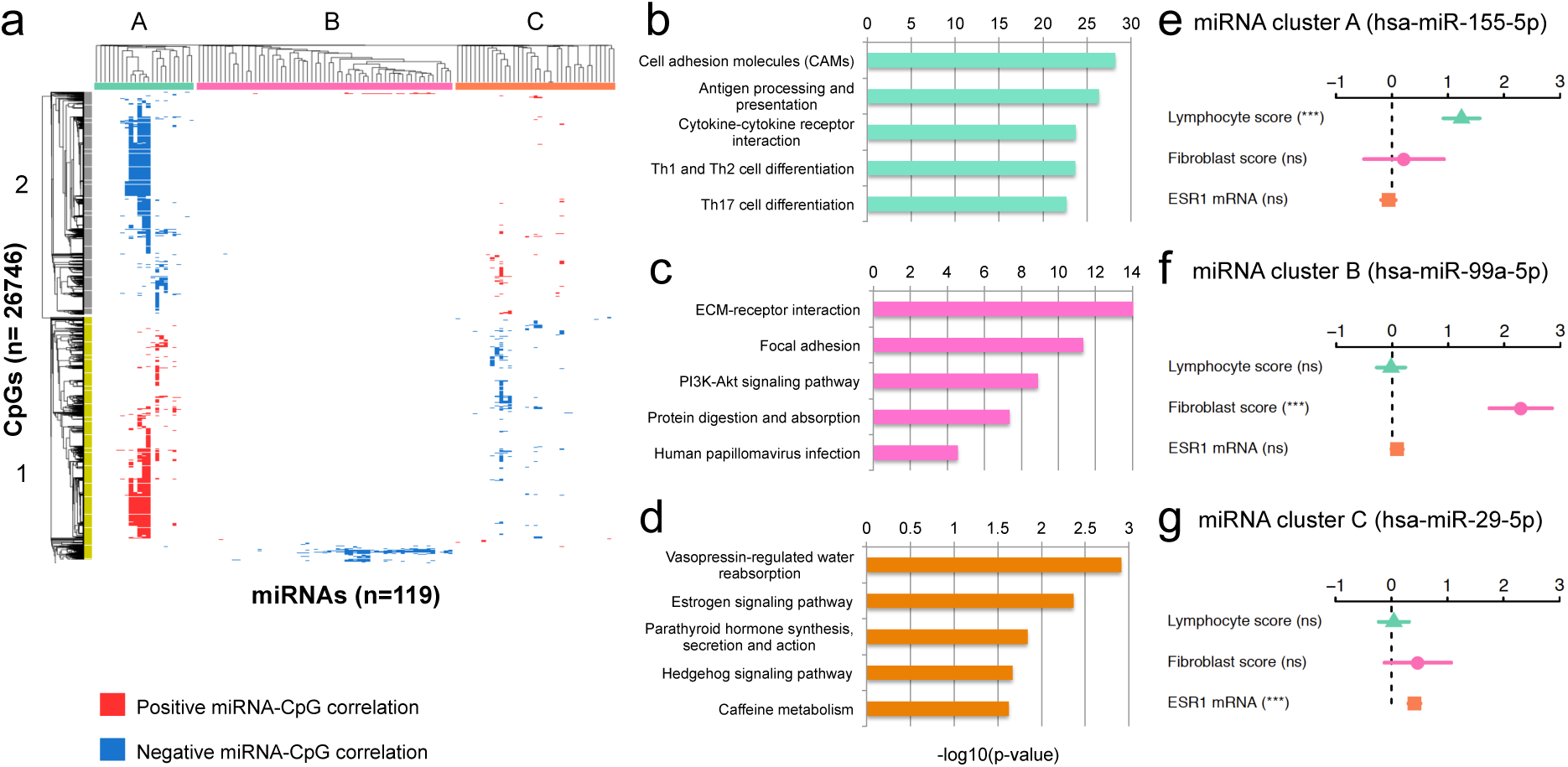
Identification of miRNA-methylation Quantitative Trait Loci (mimQTLs) clusters and corresponding annotation. **a)** Heatmap showing hierarchical clustering of the 89 118 significant mimQTLs found in both the Oslo2 and TCGA cohorts. miRNAs are shown in columns and CpGs in rows with blue color indicating a negative correlation and red color indicating a positive correlation between miRNA expression and CpG methylation. Three main miRNA clusters (cluster A, B and C) and two main CpG clusters (cluster 1 and 2) were identified. **b-d)** Barplots showing the top five most enriched pathways for genes co-expressed (miRNA-mRNA expression Spearman correlation >0.4) with the miRNAs of cluster A (**b**), B (**c**) and C (**d**). The x-axes show the –log10(p-value) of the pathway enrichment obtained from Enrichr [22]. Bars are color-coded according to the associated miRNA cluster. **e-g)** Results from fitting generalized linear models (GLM) to model miRNA expression as a multivariate function of lymphocyte infiltration (obtained by Nanodissect [23]), fibroblast infiltration (obtained by xCell [24]) and ESR1 mRNA expression. The GLM coefficients are depicted with 95% confidence intervals for each of the top miRNAs from each miRNA cluster. Asterisks (***) denote a p-value<0.001 and “ns” denotes *not significant* (p-value >0.05).

### miRNA clusters highlight important processes of breast cancer pathogenesis

To identify biological functions shared by miRNAs in the same cluster (x-axis of the heatmap Figure 1), we identified genes positively co-expressed with the miRNAs of each cluster (mRNA-miRNA Spearman correlation >0.4 in both cohorts; Additional file 4c) and performed gene set enrichment analyses (GSEA) using Enrichr [22].

#### miRNA cluster A – the immune cluster

miRNAs in cluster A were co-expressed with genes involved in immune cell differentiation and signaling (Figure 1b and Additional file 4d). The top five miRNAs with most correlations to CpGs in Cluster A (and also overall) were hsa-miR-155-5p (n=14 469), hsa-miR-146a-5p (n=12 546), hsa-miR-150-5p (n=11 679), hsa-miR-142-5p (n=8320), and hsa-miR-135b-5p (n=5766). Concordant with the GSEA, we have previously shown in a third independent breast cancer cohort that these miRNAs are highly associated with immune response processes [10]. To further confirm the association between miRNA cluster A and immune response, we used gene expression to score lymphocyte infiltration in each tumor using Nanodissect [23]. We found an enrichment of positive correlations between cluster A miRNA expression and tumor immune infiltration in both cohorts (hypergeometric test p-value<0.001 considering the correlation between all miRNAs and the lymphocyte score as background; Additional file 8a, b and Additional file 4e). Altogether, these results suggest that miRNAs in cluster A are either expressed by tumor infiltrating immune cells or mold the tumor microenvironment. This is further supported by their higher expression in estrogen receptor (ER) negative tumors (Additional file 4f and Additional file 9a, b), which have higher immune infiltration compared to ER positive tumors [25, 26].

#### miRNA cluster B – the fibroblast cluster

miRNAs in cluster B were co-expressed with genes enriched for extracellular matrix (ECM) and focal adhesion (Figure 1c and Additional file 4d). As fibroblasts are strongly associated with biophysical forces of the tumor microenvironment and in shaping the ECM through the deposition of collagen [27], we computed a score reflecting the relative amount of fibroblasts in each sample using gene expression and the xCell [24] algorithm. We found that the expression of miRNAs in cluster B was significantly enriched for positive correlations to the fibroblast score (hypergeometric test p-value<0.001 considering the correlation between all miRNAs and the fibroblast score as background; Additional file 8c, d and Additional file 4e). miRNAs of this cluster showed in general higher expression in ER positive compared to ER negative ones in the Oslo2 cohort, and consistent differential expression between PAM50 subtypes with highest expression found in the luminal A and normal-like subtypes (Additional file 4f and Additional file 9c, d).

#### miRNA cluster C – the estrogen signaling cluster

miRNAs in cluster C were co-expressed with genes associated with hormone-regulated processes lead by estrogen signaling (Figure 1d and Additional file 4d). Indeed, in both Oslo2 and TCGA the expression of cluster C miRNAs was significantly enriched for positive correlations to estrogen receptor mRNA (ESR1) expression (hypergeometric test p-value<0.001 considering the correlation between all miRNAs and ESR1 mRNA as background; Additional file 8e, f and Additional file 4e), and the miRNAs were mostly upregulated in ER positive tumors compared to ER negative ones (Additional file 4f and Additional file 9e, f). We identified hsa-miR-29c-5p as the hub of cluster C with the highest number of associations to CpG methylation (n=4764). This miRNA has been previously identified as one of the most significantly differentially expressed miRNAs between ER positive and negative tumors [10]. Thus, while miRNAs in cluster A and B reflected heterogeneity within the tumor microenvironment, miRNAs in cluster C were associated with estrogen signaling and ER positive versus negative breast cancer diseases.

To further investigate the association between miRNA expression and the three characteristics of the clusters identified above (immune and fibroblast infiltration, and ER status), we modeled miRNA expression as a multivariate function of lymphocyte and fibroblast infiltration as well as ESR1 mRNA expression. Figure 1e-g shows the coefficients for each characteristic to predict the expression of the miRNA with the highest number of CpG associations in each cluster. For nine out of 23 miRNAs in cluster A (Additional file 4g), including hsa-miR-155-5p, the ‘hub’ of cluster A (Figure 1e), the lymphocyte infiltration score was the most significant explanatory variable for expression. In cluster B, the fibroblast infiltration score was the variable significantly associated with miRNA expression for 51 out of 59 miRNAs across both cohorts as demonstrated for hsa-miR-99a-5p (Figure 1f and Additional file 4g). For cluster C (Figure 1g), ESR1 mRNA expression (surrogate for ER status) was significantly associated with hsa-miR-29c-5p expression, the hub of cluster C. Altogether, our analysis of miRNA expression in the three mimQTL-miRNA clusters clearly identified distinct signaling pathways and processes associated with different biological and molecular aspects of breast cancer.

### CpGs in mimQTL clusters reside in chromatin contexts associated with breast cancer subtypes

Next, we aimed to biologically annotate the CpGs of clusters 1 and 2, starting with their genomic position. First, we assessed pathway enrichment of their closest associated gene to infer any functional pathway association. Second, ChromHMM segmentation of the genome of several cell lines spanning breast cancer subtypes and ATAC-seq data was analyzed to study the genomic context of the CpGs within cluster. Finally, we assessed their overlap with transcription factor (TF) binding sites (TFBSs) derived from computational TF binding models and Chromatin Immunoprecipitation Sequencing (ChIP-seq) data [28].

#### Cluster 1 CpGs

CpGs in cluster 1 mapped to 4809 genes according to the annotation of the Illumina HumanMethylation450k array. With GSEA using Enrichr [22], we found these genes enriched in signaling and cancer-associated pathways such as the Ras and PI3K-Akt signaling pathways (Additional file 4h). According to the ChromHMM genome segmentation of breast cancer cell lines [29], cluster 1 CpGs were enriched at enhancers, especially of ER positive/luminal cell lines (Figure 2a and Additional file 4i). ATAC-seq data from TCGA confirmed that the regions surrounding these CpGs were more accessible (open) in ER positive than in ER negative tumors (Wilcoxon p-value = 6.38e-5; Figure 2b). Furthermore, using the UniBind [28] database storing direct TF-DNA interactions for 231 TFs using 1983 human ChIP-seq data sets, we found cluster 1 CpGs enriched at FOXA1/2, GATA2/3, TFAP2C, and ESR1 (encoding ER-alpha) binding sites; these TFs are known to drive ER positive breast cancers [2, 30] (Figure 2c). Unsupervised clustering of the DNA methylation values associated with cluster 1 CpGs separated the tumors according to breast cancer subtypes (Figure 2d and Additional file 10a for the Oslo2 and TCGA cohorts, respectively). Cluster 1 CpGs showed overall lower DNA methylation in ER positive and luminal breast cancer subtypes (Figure 2e and Additional file 11a). These lines of evidence show that cluster 1 CpGs are found at accessible enhancers with ER-associated TFBSs and are hypomethylated in ER positive/luminal tumors.

**Figure 2.**
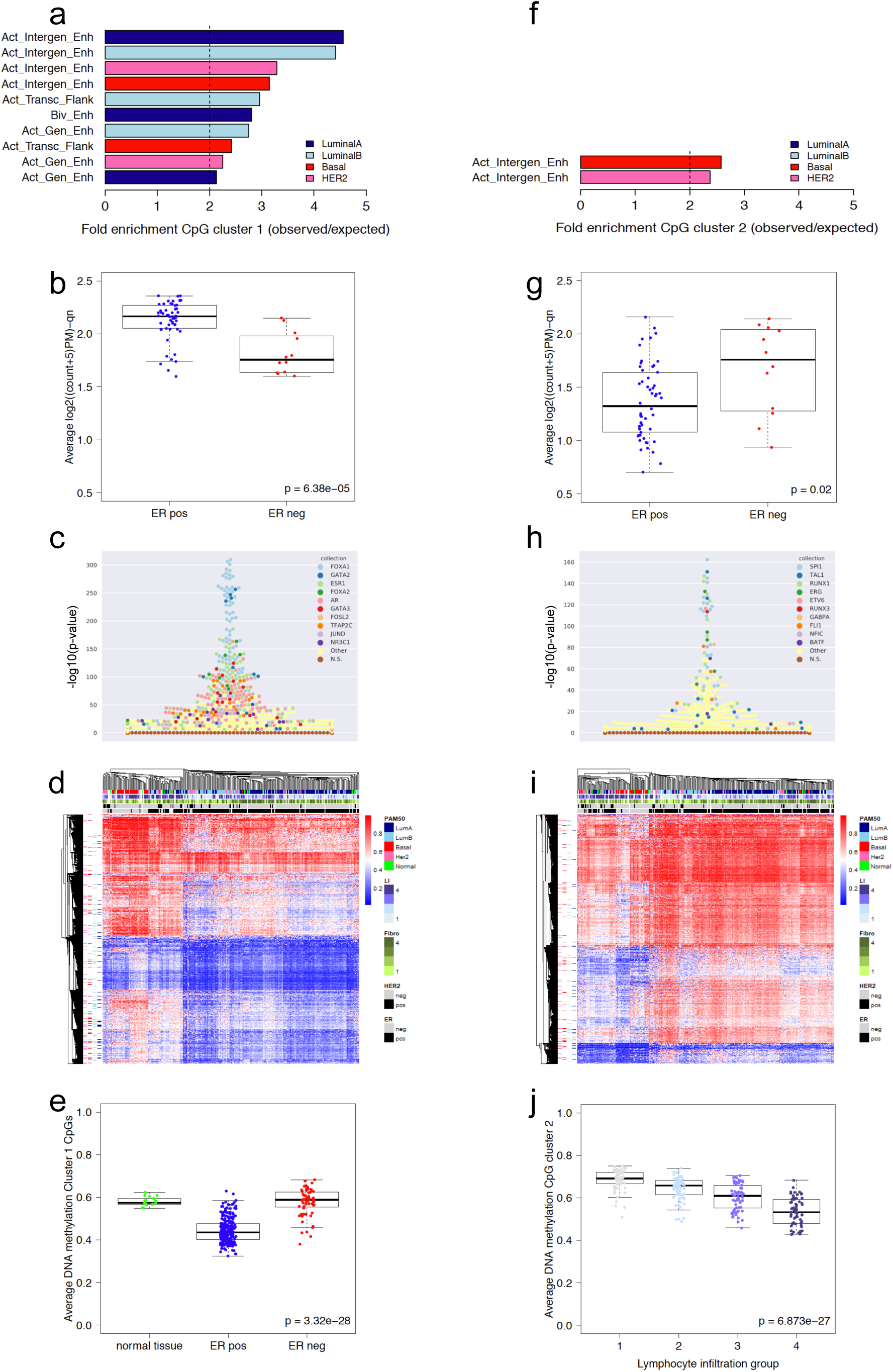
Functional annotation of the CpG clusters. **a)** Genomic location enrichment of mimQTL CpGs in cluster 1 according to ChromHMM data from cell lines representing different breast cancer subtypes [29]. Active Genic Enhancer=Act_Gen_Enh, Active Transcription Flanking=Act_Transc_Flank, Bivalent Enhancer=Biv_Enh, Active Intergenic Enhancer=Act_Intergen_Enh. **b)** Average normalized counts per tumor sample for all ATAC-seq peaks mapped to CpGs of cluster 1 (TCGA data). **c)** Swarm plot showing enrichment of TF binding sites (–(log10(p-value) using Fisher’s exact tests) on the y-axis for CpGs of cluster 1 (n=14 040) according to UniBind [28]. TF names of the top 10 enriched TF binding sites data sets are provided with dedicated colors. Data sets for the same TFs are highlighted with the corresponding colors. **d)** Heatmap showing hierarchical clustering of methylation levels of CpG cluster 1 (n=14 040) in the Oslo2 cohort (CpGs in rows and tumors in columns). Tumors are annotated according to PAM50 molecular subtypes; Lymphocyte infiltration (LI) quartile groups 1(low)–4(high); Fibroblast infiltration quartile groups (Fibro): 1(low)–4(high); Human epidermal growth factor receptor 2 (HER2) status; Estrogen receptor (ER) status. CpGs are annotated according to overlap with regions annotated as “Active Intergenic Enhancer” from ChromHMM. **e)** Boxplot showing average DNA methylation of CpGs from cluster 1 in normal breast tissue (n=17), ER positive (pos; n=223) and negative tumors (neg; n=60) of the Oslo2 cohort. **f)** Enrichment of mimQTL CpGs in cluster 2 according to ChromHMM data. **g)** Average normalized counts for ATAC-seq peaks mapped to CpGs of cluster 2. **h)** Enrichment of TF binding sites for CpGs of cluster 2 (n=12 706). **i)** Hierarchical clustering of methylation levels of CpG cluster 2 (n=12 706). **j)** Boxplot showing average DNA methylation of Cluster 2 CpGs when Oslo2 tumors were separated into lymphocyte infiltration quartile groups from low (1) to high (4). Wilcoxon rank-sum p-values (two-group comparisons) and Kruskal-Wallis p-values (three or more groups) are indicated.

#### Cluster 2 CpGs

Applying similar analyses to cluster 2 CpGs, we found the nearest genes (n=3865) associated with cancer and immune system-related pathways (Additional file 4j). Cluster 2 CpGs were enriched at breast cancer enhancer regions, but to a lower extent than cluster 1 CpGs, and more at enhancers from ER negative cell lines (Figure 2f and Additional file 4i). Further, CpGs in cluster 2 were at genomic regions more open in ER negative tumors than in ER positive ones, according to ATAC-seq data (Wilcoxon p-value = 0.02; Figure 2g). TFBSs associated with TFs involved in hematopoiesis and immune processes such as SPI1, TAL1, and RUNX1 were enriched close to cluster 2 CpGs (Figure 2h). Unsupervised clustering using the DNA methylation of cluster 2 CpGs grouped breast cancer samples according to their level of lymphocyte infiltration derived from the Nanodissect scores (Figure 2i). Finally, DNA methylation of cluster 2 CpGs negatively correlated with lymphocyte scores (i.e. low methylation – high lymphocyte infiltration and vice versa; Figure 2j and Additional file 11b). Thus, the methylation of cluster 2 CpGs is driven by intra-tumor heterogeneity characterized by infiltration of immune cells.

### Functional interpretation of CpG and miRNA mimQTL clusters

#### Functional association between CpG cluster 1 and miRNA cluster C

Altogether, we found cluster 1 CpGs to be associated with regulatory regions important for ER signaling and residing in more open and less methylated genomic regions in ER positive tumors than in ER negative ones (Figure 2a, b). Hence, the negative correlations between CpG cluster 1 and miRNA cluster C observed in Figure 1a represent functional CpG-miRNA associations of low methylation at cluster 1 CpGs (Figure 2e) correlated with higher expression of cluster C miRNAs in ER positive/luminal tumors (Additional file 4f).

#### Functional association between CpG cluster 2 and miRNA cluster A

On the other hand, cluster 2 CpGs were associated with immune infiltration and negatively correlated through mimQTL with cluster A miRNAs (Figure 1a), miRNAs themselves associated with immune infiltration [10]. We hypothesize that this observation is influenced by and reflect the variation in the presence of infiltrating immune cells in the tumors that have very different DNA methylation and miRNA expression than the cancer cells. To further support this interpretation we retrieved DNA methylation data from ER positive and negative breast cancer cell lines and from different immune cell types (Additional file 12). Focusing on the hub CpG of miRNA cluster A (the CpG with most associations to miRNAs in cluster A), a clear difference in methylation was observed between the cancer cell lines (hypermethylated) and immune cells (hypomethylated; Wilcoxon rank-sum p-value = 4.56e-12). This is consistent with ER negative/basal-like breast cancers showing higher immune infiltration compared to ER positive/luminal tumors [26], as the methylation of cluster 2 CpGs was significantly lower (Additional file 11c, d) and the miRNAs of cluster A were more expressed (Additional file 4f) in ER negative/basal-like tumors.

### Regulatory link between DNA methylation and miRNA expression

We next focused on the regulatory networks of miRNAs in cluster C as our mimQTL analysis highlighted regulatory regions (e.g. enhancers) linked to these miRNAs in a breast cancer subtype-specific manner. To find regulatory regions for miRNAs affected by DNA methylation, we targeted our analyses on the miRNA-CpG associations overlapping with (i) a catalogue of super-enhancers (SE) recently identified by Suzuki et al. [31] to drive miRNA expression, (ii) experimentally-derived long-range interactions with Pol2 binding (ChIA-PET Pol2 data) in the luminal MCF7 cell line [32] and (iii) binding regions for TFs known to drive ER positive breast cancers (ESR1, FOXA1 and GATA3) in the MCF7 cell line (ChIP-seq data).

Altogether, 273 mimQTLs were identified where the CpG overlaps an annotated breast miRNA SE (Additional file 4k). Interestingly, CpGs of cluster 1 were found to be enriched for residing within miRNA SEs compared to the background of all CpGs (hypergeometric test p-value= 1.29 x10^−26^), further confirming the enrichment of cluster 1 CpGs at distal regulatory regions regulating miRNA expression. Of the 273 mimQTLs, 50 represented a direct *in cis* association where the CpG was found in a miRNA SE mapping [31] with the corresponding mimQTL miRNA. These 50 *in cis* mimQTLs represented eight unique miRNAs all found to be cluster C miRNAs and all with significant negative correlations with the corresponding CpGs (Spearman correlation ranging from −0.30 to −0.68; Additional file 4k). This analysis suggests, for the first time to our knowledge, that DNA methylation at super-enhancers is an important regulatory feature for miRNA expression in breast cancer.

*In cis* mimQTLs (i.e. CpG and miRNA on the same chromosome) were enriched at long-range chromatin interactions with Pol2 as defined with ChIA-PET in the luminal MCF7 cell line [32] (hypergeometric test p-value 4.51 × 10^−4^). Altogether, 168 Pol2 ChIA-PET loops linked 120 pairs of CpGs and miRNAs in mimQTLs (Additional file 4l). These mimQTLs represented 24 unique miRNAs, of which 15 were from miRNA cluster C. Thus, with the mimQTL analysis we identified how DNA methylation at regulatory regions including super-enhancers and long-range distance interactions regulate the expression of miRNAs predominantly from miRNA cluster C.

Two examples of overlap between ChiA-PET Pol2 loops, miRNA-SE, and mimQTLs are shown for hsa-miR-342-3p/5p (Figure 3a) and hsa-let7b-5p (Figure 3b). We observed for these two examples that the SE/long-range interactions also overlap with TF binding regions for ERα, FOXA1, or GATA3. The combination of these evidences suggests that through the mimQTL analysis we identify, for miRNA cluster C, hypomethylated regulatory regions in ER positive breast cancer exemplified by hsa-miR-342-3p/5p (Figure 3c) and hsa-let7b-5p (Figure 3d), which may facilitate the binding of ERα, FOXA1, and GATA3. This may further lead to the increased expression of hsa-miR-342-3p/5p (Figure 3e) and hsa-let7b-5p (Figure 3f) in ER positive breast tumors. Altogether, these results suggest a direct regulatory link between the expression of cluster C miRNAs and DNA methylation at CpGs of ER-associated TF binding regions.

**Figure 3.**
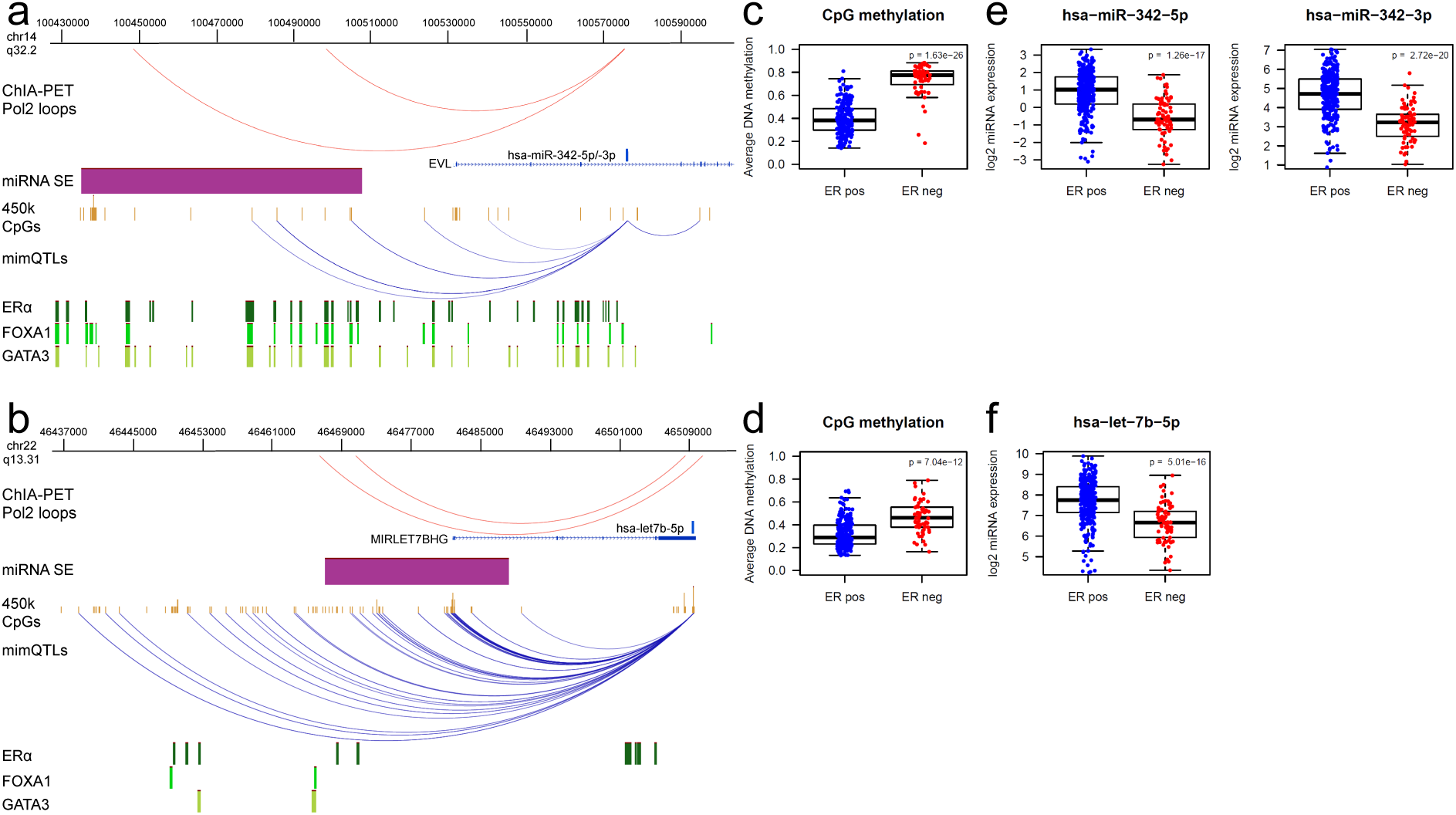
Super-enhancer (SE) - miRNA interactions and impact of CpG methylation on miRNA expression. **a)** Example of mimQTLs (blue arcs) and ChIA-PET Pol2 loops (red arcs) where mimQTL CpGs (n=3) or one foot of the ChIA-PET Pol2 loop are located within the hsa-miR-342 SE (purple) and the other loop foot reside within hsa-miR-342-5p/-3p. Also shown are the location of 450k methylation array CpGs and ERα, FOXA1 and GATA3 binding regions obtained from ChIP-Seq experiments of the MCF7 cell line. The figure was made using the WashU Epigenome Browser v. 46.2 [33]. **b)** mimQTLs and ChIA-PET Pol2 loops where mimQTL CpGs (n=21) or one foot of the ChIA-PET Pol2 loop are located within the let-7b SE (purple) and the other loop foot reside within hsa-let-7b-5p. **c)** Boxplot showing average DNA methylation in Oslo2 estrogen receptor (ER) positive (pos) and negative (neg) tumors across all CpGs within the hsa-miR-342 SE and in mimQTL with hsa-miR-342-3p/-5p (n=3). **d)** Boxplot showing average DNA methylation in Oslo2 estrogen receptor (ER) positive and negative tumors across all CpGs within the let-7b SE and in mimQTL with hsa-let-7b-5p (n=21). **e)** Boxplots showing hsa-miR-342-5p/-3p expression in ER positive and negative tumors of the Oslo2 cohort. **f)** Boxplots showing hsa-let-7b-5p expression in ER positive and negative tumors of the Oslo2 cohort. P-values resulting from Wilcoxon rank-sum tests are indicated in the boxplots.

### miRNAs associated with global breast cancer DNA methylation alterations

Further, we sought to identify miRNAs associated with methylation deregulation in breast cancer. We developed a global methylation alteration (GMA) score that reflects how overall DNA methylation in breast tumors deviates from healthy breast tissues. In brief, for each tumor, the deviation in methylation per CpG relative to that of normal breast tissue was summed up (see Methods for details). We found that ER positive and luminal B tumors showed a higher GMA score than ER negative tumors and other PAM50 subtypes (Figure 4 a-d). To identify which miRNAs may be the most potent at driving DNA methylation alterations, we correlated the expression of each of the 119 mimQTL miRNAs to the GMA score (Additional file 4m). We observed that miRNAs in cluster C were enriched for positive correlations to the GMA score when compared to the background of all miRNAs tested (hypergeometric test p-values<0.001; Figure 4 e, f). Table 1 lists the miRNAs positively correlated with the GMA score across both cohorts. Inversely, cluster A and B miRNAs were enriched for negative correlations with the GMA score (hypergeometric test p-values<0.001). Such cluster-specific associations to the GMA score was further illustrated by plotting each miRNA’s expression correlation with the GMA score according to clusters (Figure 4 g, h).

**Table 1.**
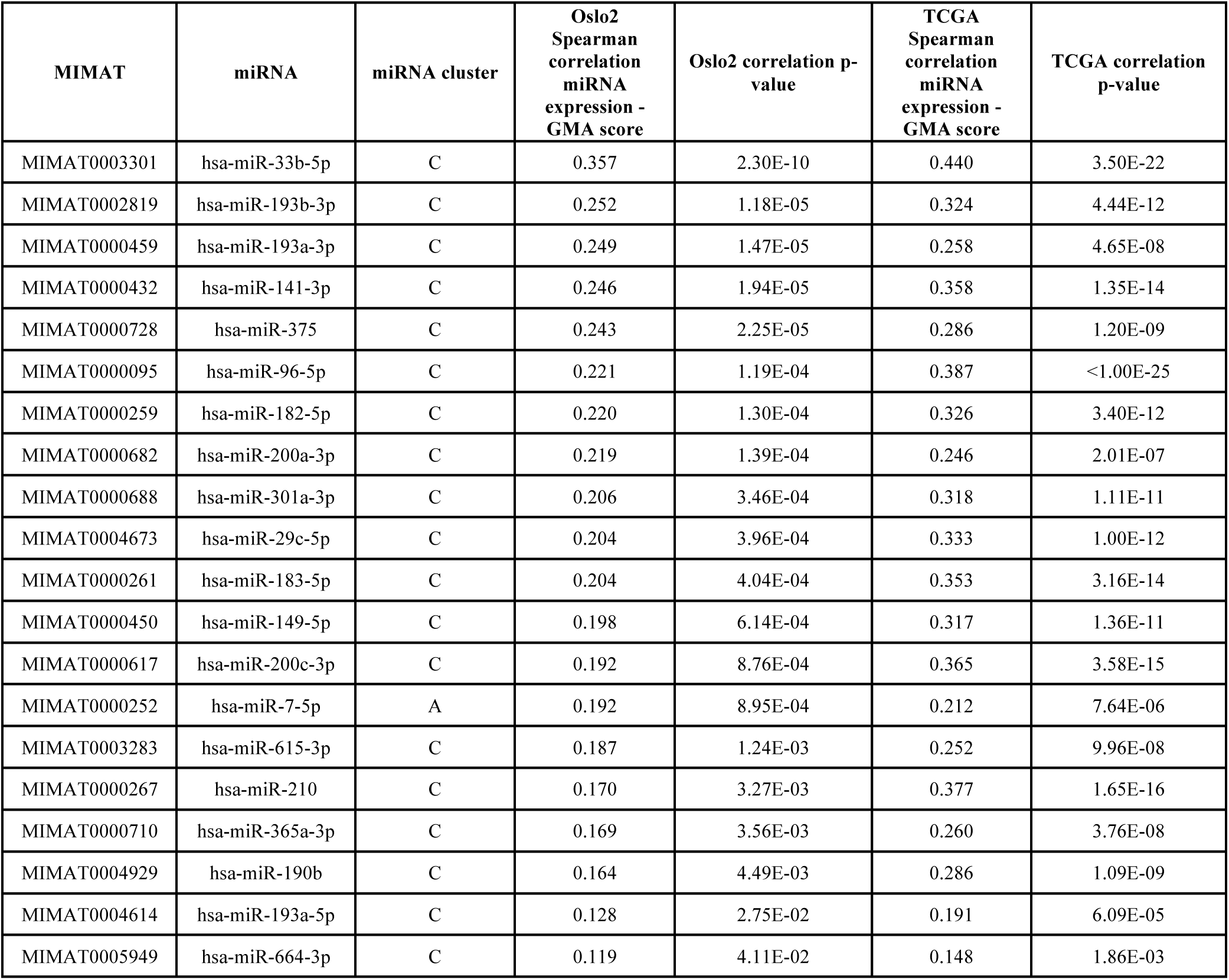
miRNAs significantly positively correlated with the global methylation alteration (GMA) score across both cohorts. The table is sorted according to correlations in the Oslo2 cohort.

**Figure 4.**
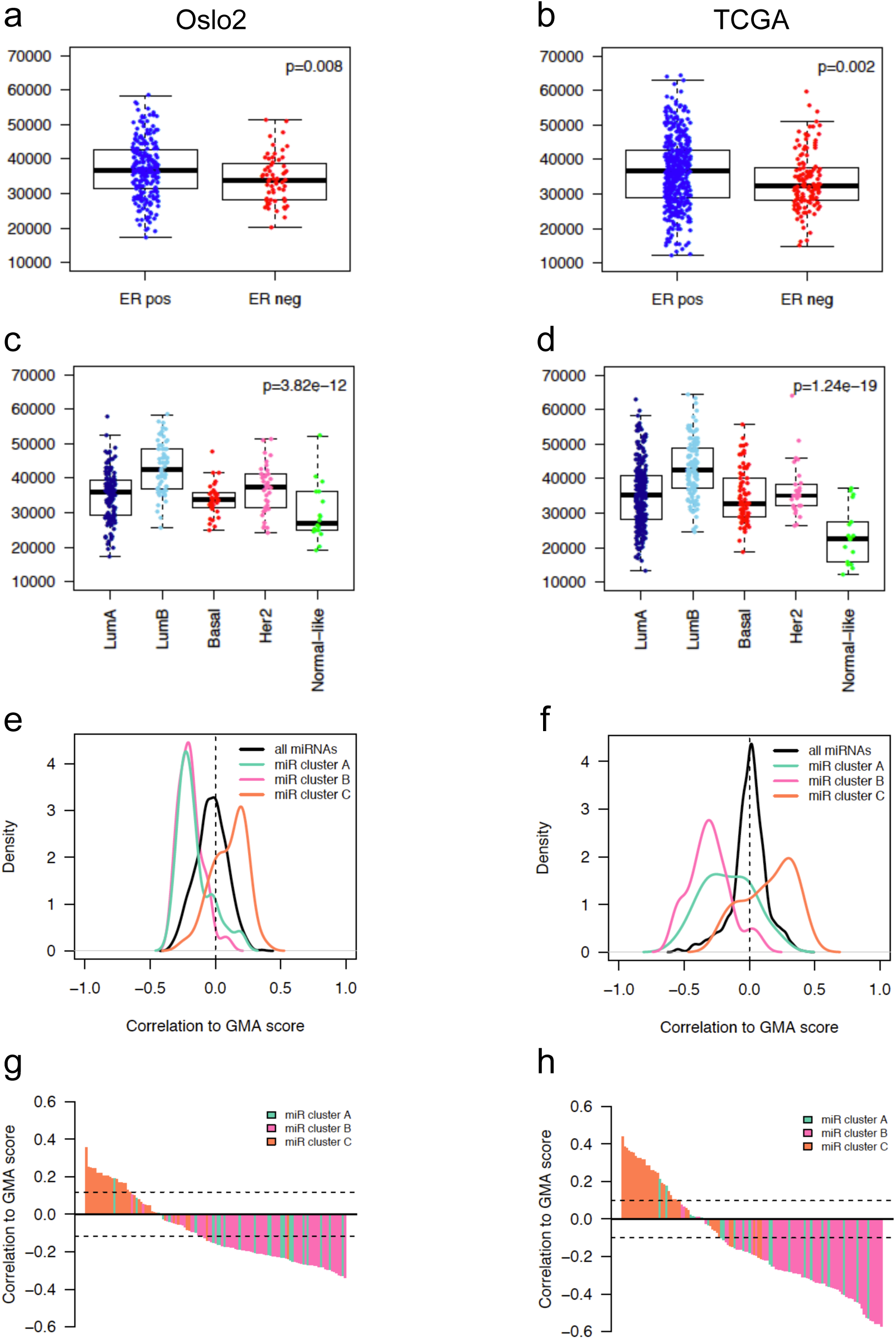
Global Methylation Alteration (GMA) score in clinical breast cancer groups and correlation to miRNA expression. **a, b)** Boxplots showing the GMA score in estrogen receptor (ER) positive (pos) and negative (neg) tumors of the Oslo2 cohort **(a)** and TCGA cohorts **(b)**. Wilcoxon rank-sum test p-values are denoted. **c, d)** Boxplots showing the GMA score in PAM50 molecular subtypes of the Oslo2 **(c)** and TCGA cohorts **(d)**. LumA: Luminal A, LumB: Luminal B, Basal: Basal-like, Her2: HER2-enriched. Kruskal-Wallis test p-values are denoted. **e, f)** Plots showing density curves of the correlation between miRNA cluster members and the GMA score for the Oslo2 **(e)** and TCGA **(f)** cohorts. The density lines are color-coded according to miRNA cluster. **g, h)** Barplots showing miRNAs decreasingly ranked according to GMA score correlation level (y-axis) in the Oslo2 **(g)** and TCGA cohorts **(h)**. The bars are color-coded according to miRNA cluster.

### hsa-miR-29c-5p is negatively correlated to DNA methyltransferase 3A and is deregulated early during breast cancer pathogenesis

Hypothesizing that cluster C miRNAs may be positively correlated with the GMA score through regulation of enzymes involved in DNA methylation, we assessed the *in silico* target prediction database TargetScan [34] focusing on enzymes regulating DNA methylation (DNMTs and TETs). Altogether, 18 unique miRNAs belonging to miRNA cluster C were predicted to target DNMTs and/or TETs (Additional file 4n). We further found that three of the 18 miRNAs showed consistent and significant negative correlation to the mRNA of DNA methylation regulating enzymes (DNMT3A - hsa-miR-29c-5p (Figure 5a), TET1 - hsa-miR-365a-3p and TET1 - hsa-miR-375; Additional file 4n). We further confirmed the negative correlation between hsa-miR-29c-5p and DNMT3A also at the protein level (Figure 5b, Spearman’s rho=-0.77) using proteome data [35] of the Oslo2 samples (n=45). Note that this inverse correlation between hsa-miR-29c-5p and DNMT3A protein levels was the third most negative correlation considering all correlations between miRNAs (n=713) and proteins (n=9995) in the Oslo2 cohort. Of note, a significant negative correlation was also observed between hsa-miR-29c-5p and DNMT3B/DNMT1 mRNA and protein levels (Additional file 13).

**Figure 5.**
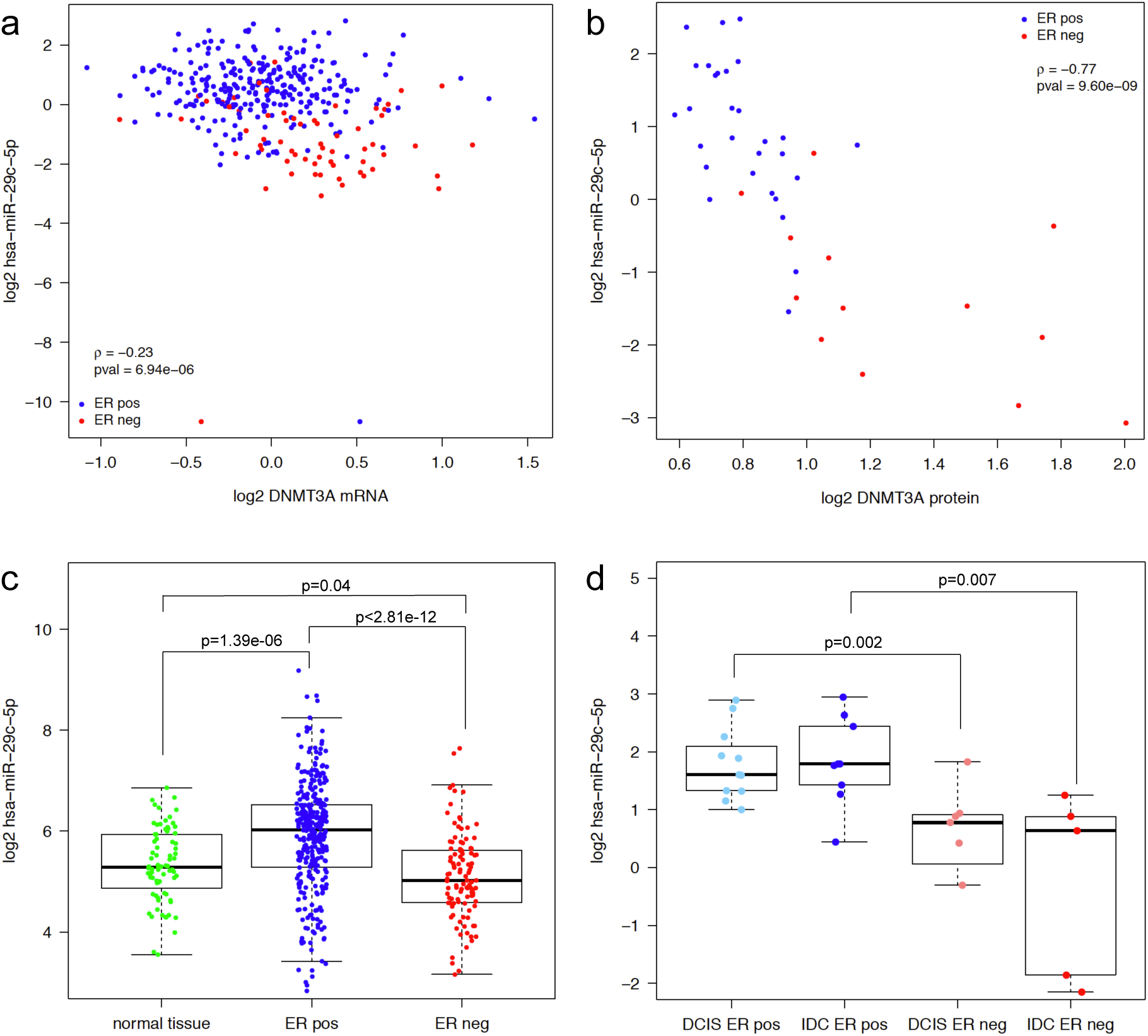
Expression of hsa-miR-29c-5p and correlation to DNMT3A. **a)** *DNMT3A* mRNA expression (x-axis) vs. hsa-miR-29c-5p expression (y-axis) measured in 377 samples of the Oslo2 cohort. Estrogen receptor (ER) positive (pos) tumors are plotted in blue and ER negative (neg) in red. **b)** DNMT3A protein expression (x-axis) vs. hsa-miR-29c-5p expression (y-axis) measured in 45 samples of the Oslo2 cohort. **c)** hsa-miR-29c-5p expression in normal adjacent breast tissue (normal; n=76), ER positive (n=333) and ER negative tumors (n=106) of the TCGA cohort. Wilcoxon rank-sum test p-values are denoted. **d)** hsa-miR-29c-5p expression in ER positive (n=11) and ER negative (n=7) ductal carcinoma *in situ* (DCIS) samples and ER positive (n=9) and ER negative (n=5) invasive ductal carcinoma (IDC) samples from the same data set [13]. Wilcoxon rank-sum test p-values are denoted.

Furthermore, hsa-miR-29c-5p was found significantly upregulated in ER positive tumors compared to normal breast tissue and ER negative tumors (Wilcoxon rank-sum p-value = 1.39 × 10^−6^ and p-value < 2.81 × 10^−12^; Figure 5c). To investigate whether the changes of hsa-miR-29c-5p expression may happen early during breast carcinogenesis causing DNA methylation alterations, we mined ductal carcinoma *in situ* (DCIS) samples (n=18) from an independent data set [13]. Indeed, in ER positive pre-invasive DCIS lesions, hsa-miR-29c-5p was significantly higher expressed than in ER negative DCIS lesions (Wilcoxon rank-sum p-value=0.002; Figure 5d), indicating that initial changes in hsa-miR-29c-5p expression may drive early DNA methylation changes in DCIS as previously observed [1].

## Discussion

This study is to our knowledge the first to assess the global relationship between the expression of miRNAs and DNA methylation on a genome-wide scale in breast cancer. Using two large and independent breast cancer cohorts we have identified robust associations that point to how miRNA expression may be regulated through methylation at distal regulatory regions and how miRNAs may contribute to mold the epigenetic landscape of breast cancer. Further, the analysis points to how miRNA expression may reflect levels of infiltration from the surrounding microenvironment. The mimQTL approach identifies a positive or negative sign of the relation between a miRNA and a CpG, but does not infer the *causality* of the association, if any. Grouping of the mimQTLs using hierarchical clustering revealed aspects of underlying intra- and inter-tumor heterogeneity in the form of immune cell or fibroblast infiltration and tumor ER status. Expression of miRNAs in cluster A were positively associated with the lymphocyte score reflecting immune infiltration. Indeed, the miRNAs in this cluster with most CpG associations, hsa-miR-155-5p, hsa-miR-146a-5p, hsa-miR-150-5p, and hsa-miR-142-5p, have previously been associated with immune related pathways [10] and lymphocytic infiltration [16] in breast cancer. Furthermore, in a study characterizing miRNA expression in various cell types and tissues from McCall et al. [9], these miRNAs were found to be most highly expressed in B- and T-cells, further suggesting that our miRNA cluster A reflect signals coming from infiltrating immune cells. This underlies a need for understanding more about which type of cells from bulk tumor samples are actually expressing nominated miRNA cancer biomarkers. Importantly, the context- and cell type specific expression of miRNAs give them an attractive potential for deconvolution tools. Using the deconvolution tool xCell [24] based on mRNA expression, we linked miRNAs in cluster B to fibroblast cells as their expression was positively correlated with the fibroblast score. Fibroblasts are providers of ECM-components [36] and the genes co-expressed with the miRNAs were enriched for ECM-associated pathways. Assessing the catalogue of cell- and tissue type specific expression of miRNAs [9] supported this finding with the top miRNAs of this cluster, hsa-miR-99a-5p, hsa-miR-125b-5p, hsa-miR-379-5p, hsa-miR-381, and hsa-miR-100-5p, being highly expressed in tissue from skin where fibroblasts are a major component. Interestingly, hsa-miR-125b-5p was found to induce cardiac fibrosis [37]. Other studies pointed to a tumor-suppressor role of these miRNAs in breast cancer, for instance hsa-miR-99a-5p reduces breast cancer cell viability by targeting mTOR [38], hsa-miR-125b-5p was shown to induce cell cycle arrest and reduce cell growth in breast cancer cells [39] and hsa-miR-379-5p was shown to regulate Cyclin B1 expression [40]. More studies using for instance *in situ* hybridization of tumor tissue sections are needed to further validate the cells of origin of the immune- and fibroblast-associated miRNAs of cluster A and B, respectively, and will help to further refine the role of these miRNAs in breast cancer.

The miRNAs with most CpG associations in cluster C were markers of the ER positive, luminal phenotype of breast cancer [10]. Integrating mimQTLs with data from various sources including ChIA-PET Pol2 loops, ATAC-seq, ChIP-seq, and miRNA SEs we showed how their expression may be promoted by ensuring open and active enhancer regions where ER-associated TFs bind and loop to the miRNA-encoding genomic regions boosting both their transcription and processing [31]. We speculate that the observed demethylation of the miRNA SE CpGs in ER positive tumors lead to binding of ER-associated TFs making the SE active. Through activation, the SE is looped with Drosha/DGCR8 - a protein complex important for processing of the primary miRNA transcript to the shorter precursor transcript [41]. This SE-mediated miRNA processing was previously shown with ChIP-seq peaks for DGCR8 observed at both the transcription start site (TSS) and precursor miRNA regions for SE-associated miRNAs [31]. The looping of the miRNA SE to the miRNA TSS or mature sequence boost the transcription and the processing of the miRNA [31], which was in our study further supported by the negative mimQTL correlation; low SE CpG methylation associated with high miRNA expression in ER positive tumors. This emphasizes an important regulatory role for SE CpG methylation on miRNA expression in breast cancer. Indeed, miRNA expression deregulation in breast cancer through methylation alterations was previously described [16, 21, 42], but the focus has mainly been on CpGs in proximal promoter regions. As the importance of enhancer region conformation and methylation is becoming increasingly appreciated and given the great impact of miRNAs on the establishment and maintenance of cell phenotype, exploring this field will give new insights into cancer development and progression. Cluster C miRNAs were consistently enriched for positive correlations to the GMA score indicating that ER positive, luminal tumors may be more severely altered at the methylation level compared to ER negative tumors which may be more driven by alterations at the copy number level [12] and which showed methylation patterns more similar to the normal breast tissue samples.

Our analyses of the genomic positions and methylation of CpGs in each cluster highlighted CpG cluster 1 associated with differences in DNA methylation of enhancers and TFBS according to ER status. On the other hand, cluster 2 was associated with intra-tumor heterogeneity and infiltration of immune cells. Importantly, our CpG analyses mapped back to the biological functions associated with the miRNA clusters and therefore point to the fact that not only correlative but also functional associations link (i) CpG-cluster 1 and miRNA cluster C as both being associated with estrogen response and ER status and (ii) CpG cluster 2 and miRNA cluster A being related to tumor immune infiltration. However, it is important to note that inter-tumor heterogeneity defined by ER status and intra-tumor heterogeneity defined by immune infiltration are to some aspects two sides of the same coin as ER negative tumors show a higher degree of immune infiltration [26].

We identified hsa-miR-29c-5p as a potential epigenetic hub in ER positive breast cancer as it was the miRNA in cluster C with most CpG associations, positively correlated with the GMA score, upregulated in ER positive tumors compared to both ER negative tumors and normal breast tissue, *in silico* predicted to target DNMT3A and negatively correlated to DNMT3A mRNA and protein levels. The miR-29 family has previously been shown to directly target DNMTs [19], confirming a role for these miRNAs as epigenetic regulators, although for hsa-miR-29c the attention has been on the −3p mature form rather than the −5p. Further functional validation is required to prove the direct causal role of hsa-miR-29c-5p in determining luminal breast cancer phenotype. Supporting our hypothesis of hsa-miR-29c-5p being important for establishing the ER positive/luminal breast cancer phenotype by targeting DNMT3A which further leads to hypomethylation of CpGs at ER-associated TFBSs, Chou et al. [43] found that GATA3 acts as a TF inducing the expression of the miR-29 family. This and other studies have, however, pointed to a tumor-suppressor role of the miR-29 family in breast cancer as they are typically found higher expressed in less aggressive/better prognosis subtypes and with over-expression in cell lines inhibiting metastasis, proliferation, migration and growth [43-45]. Nevertheless, these findings are not contradictory with our hypothesis of the epigenetic regulator role of hsa-miR-29c-5p within luminal phenotypes. Importantly, focusing on subtype-specific progression we previously found in an independent data set that hsa-miR-29c-5p is upregulated in expression from DCIS to luminal A and B tumors supporting the potential role of this miRNA in breast cancer progression within ER positive tumors [14].

In conclusion, we find that CpG methylation at ER-associated TF binding regions is likely to be important for regulation of miRNA expression in breast cancer. Furthermore, our study highlights that deregulation of hsa-miR-29c-5p expression is an early event that may result in downregulation of DNMT3A, which could further lead to hypomethylation of CpG sites important for ER positive breast cancer cell identity. The CpG sites affected are at enhancer regions with TFBS for ER-alpha, FOXA1, and GATA3, all known to be important for the luminal breast cancer phenotype.

## Methods

### Clinical materials

Two independent breast cancer cohorts with DNA methylation and miRNA expression available were here used in parallel; the Oslo2 breast cancer cohort [17, 18] and The Cancer Genome Atlas Breast Invasive Carcinoma (TCGA-BRCA) cohort [12].

The Oslo2 breast cancer cohort has been previously described [17, 18] and is a consecutive study collecting material from breast cancer patients with primary operable disease at several hospitals in south-eastern Norway. Patients were included in the years 2006-2019. The study was approved by the Norwegian Regional Committee for Medical Research Ethics (approval number 1.2006.1607, amendment 1.2007.1125), and patients have given written consent for the use of material for research purposes. All experimental methods performed are in compliance with the Helsinki Declaration.

The Illumina Infinium HumanMethylation450 microarray was used to measure the DNA methylation levels of more than 450 000 CpG sites for 330 patient tumors from the Oslo2 cohort as previously described [1, 46]. Preprocessing and normalization involved steps of probe filtering, color bias correction, background subtraction and subset quantile normalization. The DNA methylation data have been previously published [2]. For comparison of Oslo2 CpG DNA methylation levels to normal tissue, data from normal breast tissue from reduction mammoplasty (n=17) were available [1].

The one-color microarray Human miRNA Microarray Kit (V2) design ID 029297 from Agilent Technologies was used to measure miRNA expression for 425 tumors of the Oslo2 cohort using 100 ng total RNA as input. Scanning was performed on the Agilent Scanner G2565A. Samples were processed using Feature Extraction version 10.7.3.1 (Agilent Technologies). The data were log_2_-transformed and for each tumor sample, considering only expressed miRNAs, the data were median centered. All non-expressed miRNAs across tumors were set to a common minimum value. The miRNA and mRNA expression data have been previously published [17]. In total, 297 Oslo2 tumor samples had matched methylation and miRNA expression data. Furthermore, of these, 45 samples had protein expression available measured by mass spectrometry and published in Johansson et al. [35].

The Cancer Genome Atlas Breast Invasive Carcinoma (TCGA-BRCA) cohort [12], from here on named TCGA, has been previously described [12]. For the DNA methylation data (level 3), probes with more than 50% missing values were removed, and further missing values were imputed using the function pamr.knnimpute (R package *pamr*) with k = 10. The log2(RPM+1) miRNA mature strand expression data (level 3) measured by IlluminaHiseq were downloaded from the UCSC Xena browser [47]. In case of NAs, these were replaced with 0. Altogether, 439 TCGA breast cancer samples had matched methylation and miRNA expression data. In addition, DNA methylation level and miRNA expression data were available for 97 and 76 adjacent TCGA normal tissue samples, respectively. TCGA gene expression data in the form of log2(norm_count+1) measured by IlluminaHiseq_RNASeqV2 were downloaded from the UCSC Xena browser [47].

miRNA expression from DCIS samples were available from Lesurf et al. [13]. In this data set, 26 DCIS samples and 14 invasive ductal carcinoma (IDC) had miRNA expression data, and out of these, 18 and 14, respectively, had estrogen receptor status available.

### Statistical and bioinformatical analysis

All analyses were performed in the R software v. 3.5.3 [48] unless otherwise specified. mimQTL analysis R code is available from GitHub: https://github.com/miriamragle/mimQTL.git.

### Genome-wide correlation analysis

Within each data set, CpGs with an interquartile range (IQR) >0.1 and miRNAs expressed in >10% of the samples were selected. Considering only CpGs and miRNAs present in both data sets resulted in 142 804 CpGs and 346 miRNAs (Additional file 1). To test the correlation between the level of DNA methylation of CpGs and miRNA expression, the Spearman correlation statistics was applied (function *cor.test* with *method=“spearman”* in R). An association was considered statistically significant if a Bonferroni-corrected p-value was <0.05. Only significant correlations with the same direction (sign) were kept.

### Hierarchical clustering of mimQTLs

The significant correlations overlapping the two data sets from the genome-wide correlation analysis were transformed into binary terms with −1 for a significant negative correlation and +1 for a significant positive correlation. The hierarchical clustering of CpGs and miRNAs was performed on these values using the R package *pheatmap* version 1.0.12 [49] with correlation distance and average linkage. CpGs and miRNAs with at least one significant association were included in the clustering analysis. To identify and decide upon the number of CpG and miRNA clusters, the dendrograms were visually inspected using different cut-offs on the *cutree_rows* and *cutree_cols* functions of the *pheatmap* package. Cut-offs were manually selected to define the clusters depicted in Figure 1a (with cutree_rows = 2 and cutree_cols = 3).

### Biological annotation of miRNA clusters

For each miRNA in a given cluster, its expression was correlated to the mRNA expression of all genes. Genes that were positively correlated (Spearman correlation >0.4 and Spearman correlation p-value<0.05) in both the Oslo2 and TCGA cohorts were kept. For each miRNA cluster, the list of positively correlated genes (i.e. co-expressed genes) were selected and used as input to Enrichr [22] to perform gene set enrichment analysis (on 13.09.2019). Results obtained from the KEGG 2019 Human Pathways database were used.

### Lymphocyte and fibroblast infiltration scores

The Nanodissect algorithm [23] (http://nano.princeton.edu/) was used for *in silico* estimation of lymphocyte infiltration as previously described [2]. The xCell algorithm [24] was used to obtain a fibroblast score for Oslo2 samples. For TCGA, xCell scores were downloaded from https://xcell.ucsf.edu/xCell_TCGA_RSEM.txt. To assess enrichment of positive or negative correlations between miRNA expression and infiltration scores for a given miRNA cluster, the *phyper* function in R was used with all miRNAs (n=346) as background.

### miRNA expression modeled with generalized linear models

Generalized linear modeling (*glm* function in R) was used to model miRNA expression as a function of lymphocyte infiltration, fibroblast infiltration, and ESR1 mRNA expression to estimate which variable(s) significantly associated with miRNA expression. The estimates plotted in Figure 1 represent the multivariate analysis estimates with their 95% confidence intervals and significance of corresponding p-values are indicated.

### Pathway enrichment of genes mapped to mimQTL CpGs

For each of the CpGs in the two mimQTL CpG clusters, the corresponding gene was obtained by intersecting the Illumina450k array annotation file. The two gene lists were used as input to Enrichr [22] to perform gene set enrichment analysis (on 11.10.2019). As output we exported the results from the KEGG 2019 Human Pathways database.

### Functional annotation of mimQTL CpGs

For functional annotation of the CpGs, we utilized the ChromHMM segmentation from Xi et al. [29] obtained from cell lines representing different breast cancer molecular subtypes [29]: MCF7 and ZR751 (luminal A), MB361 and UACC812 (luminal B), AU565 and HCC1954 (HER2) and MB469 and HCC1937 (basal). These segmentations were derived from ChIP-seq data for five histone modification marks (H3K4me3, H3K4me1, H3K27me3, H3K9me3, and H3K36me3) to predict thirteen distinct chromatin states: active promoters (PrAct) and promoter flanking regions (PrFlk), active enhancers in intergenic regions (EhAct) and genic regions (EhGen), active transcription units (TxAct) and their flanking regions (TxFlk), strong (RepPC) and weak (WkREP) repressive polycomb domains, poised bivalent promoters (PrBiv) and bivalent enhancers (EhBiv), repeats/ZNF gene clusters (RpZNF), heterochromatin (Htchr), and quiescent/low signal regions (QsLow). We assessed enrichment of CpG sets within each of the 13 chromatin states using hypergeometric tests (the R function *phyper*) with all Illumina Infinium HumanMethylation450 BeadChip CpGs as background (n=485 512). P-values were corrected using the Benjamini-Hochberg (BH) procedure [50].

Normalized ATAC-seq peak signals (log2((count+5)PM)-qn) for 74 TCGA breast tumors were downloaded from the Xena browser [47] (https://xenabrowser.net/datapages/). The CpG positions from the Illumina 450k array were intersected with the peaks using BEDTools v2.29.2 [51]. To test for differential open regions between ER positive and negative tumors, the average normalized counts of the peaks containing each CpG within a CpG cluster was calculated per tumor and a Wilcoxon rank-sum test was applied to test for statistical significance using R.

### Enrichment of mimQTL CpGs at TF binding regions

To assess the enrichment of mimQTL CpGs close to TFBSs, we considered the direct TF-DNA interactions (i.e. TFBSs) stored in the UniBind database [28] at https://unibind.uio.no. These TFBSs were obtained by combining both experimental (through ChIP-seq) and computational (through position weight matrices from JASPAR [52]) evidence of direct TF-DNA interactions (see Gheorghe *et al*. [28] for more details) for 231 TFs in 315 cell lines and tissues. Note that a TF can have multiple sets of TFBSs derived from different ChIP-seq experiments. The genomic positions of all CpGs from the Illumina 450k array were lifted over from hg19 to hg38 and extended with 100 bp on each side using BedTools (v2.26.0). The enrichment of UniBind TFBS sets in regions surrounding clusters 1 and 2 CpGs were assessed against a universe considering all CpG regions with the UniBind enrichment tool (https://unibind.uio.no/enrichment/, source code available at https://bitbucket.org/CBGR/unibind_enrichment/). Specifically, the enrichment is computed using the LOLA R package (version 1.12.0) [53] using Fisher’s exact tests. Figure 2 c, h plots the Fisher’s exact p-values using swarm plots (*swarmplot* function of the *seaborn* Python package, https://doi.org/10.5281/zenodo.824567) with annotations for the TFs associated with top 10 most enriched TFBS sets.

### Hierarchical clustering of methylation and miRNA expression

Hierarchical clustering of CpG DNA methylation or miRNA expression was performed using the R package *pheatmap* version 1.0.12 [49] with Euclidean distance and average linkage. For visualization, miRNA expression values were centered and scaled with *scale = “row”*.

### Statistical testing of methylation and miRNA expression between clinical groups

When two groups were compared Wilcoxon rank-sum tests were used considering a significance level of p<0.05. For three or more groups, Kruskal-Wallis tests were used with the same significance level. When many tests were performed simultaneously, the resulting p-values were corrected using the Benjamini-Hochberg procedure [50].

### ChIA-PET Pol2 data and ChIP-seq peaks

ChIA-PET Pol2 loop data from the MCF7 cell line was retrieved from ENCODE, accession number ENCSR000CAA [32]. We investigated overlaps between ChIA-PET Pol2 loops and *in cis* (on the same chromosome) mimQTLs. A mimQTL (CpG-miRNA pair) was considered to be in a ChIA-PET loop if the CpG and the miRNA precursor were found in two different feet of the same loop. Enrichment was calculated using hypergeometric tests (*phyper* R function) with all possible *in cis* (i.e. on the same chromosome) pairs between miRNAs and CpGs of the 450k array as background. For the specific analyses of MCF7 TF ChIP-seq data sets, we retrieved hg19 ENCODE ChIP-seq peak regions from the ReMap 2018 [54] database (ENCSR000BST.GATA3.MCF7, ERP000783.ESR1.MCF7, and GSE72249.FOXA1.MCF7).

### miRNA SE breast tissue overlap with CpGs

miRNA super-enhancers (SEs) were retrieved from Suzuki et al. [31]. Data from breast-associated cell lines HCC1954, HMEC, and MCF7 were considered. The overlap between CpG genomic positions and miRNA SEs were obtained using the *GenomicRanges* R package version 1.32.7 [55].

### Global Methylation Alteration (GMA) score

To obtain one score per tumor measuring the global methylation pattern deviation of that tumor from that of normal breast cells, a Global Methylation Alteration (GMA) score was defined. Starting with the processed methylation data, for each CpG, the median beta value of all normal breast tissue samples (n=17 for Oslo2 and n=97 for TCGA) were calculated. The GMA score of a tumor *i* was then computed as:

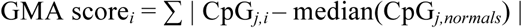

with CpG_*j,i*_ corresponding to the beta value of CpG *j* in tumor *i* and CpG_*j,normals*_ corresponding to the median of the beta values for CpG *j* in the normal breast tissue samples. Additional file 14 outlines the distribution of the GMA score in normal breast tissue samples compared to tumor samples (TCGA data) showing how tumors have a higher and much broader GMA score compared to normal breast tissue.

### *In silico* miRNA target predictions

*In silico* predicted miRNA-target interactions were downloaded from TargetScan release 7.2 [34] (http://www.targetscan.org/cgi-bin/targetscan/data_download.vert72.cgi), and both the Conserved and Nonconserved Site Context Scores were considered. From these predictions, we extracted the *Homo sapiens in silico* predictions for six selected genes: TET1, TET2, TET3, DNMT1, DNMT3A, and DNMT3B.

## Abbreviations

(ATAC-seq): Assay for Transposase-Accessible Chromatin using sequencing;
(Basal): Basal-like;
(ChIA-PET): Chromatin Interaction Analysis by Paired-End Tag Sequencing;
(ChIP-seq): Chromatin Immunoprecipitation Sequencing;
(DNMTs): DNA methyltransferases;
(DCIS): ductal carcinoma *in situ*;
(ER): estrogen receptor;
(ECM): extracellular matrix;
(GEO): Gene Expression Omnibus;
(GLM): generalized linear model;
(GMA): global methylation alteration;
(Her2): HER2-enriched;
(IQR): interquartile range;
(IDC): invasive ductal carcinoma;
(LumA): Luminal A;
(LumB): Luminal B;
(miRNA): microRNA;
(mRNA): messenger RNA;
(mimQTL): miRNA-methylation Quantitative Trait Loci;
(neg): negative;
(pos): positive;
(Pol2): RNA polymerase II;
(SE): super-enhancer;
(TCGA): The Cancer Genome Atlas;
(TCGA-BRCA): The Cancer Genome Atlas Breast Invasive Carcinoma;
(TETs): Ten-eleven translocation enzymes;
(TF): transcription factor;
(TFBS): transcription factor binding site;
(TSS): transcription start site.

## Declarations

### Ethics approval and consent to participate

Patients have given written informed consent. The Regional Committee for Medical and Health Research Ethics for southeast Norway have approved the study (approval number 1.2006.1607, amendment 1.2007.1125 for the tumor material collected at Oslo University Hospital, Norway, and approval number 429-04148 for the tumor material collected at Akershus University Hospital, Norway).

### Consent for publication

Not applicable.

### Availability of data and materials

The DNA methylation data of the Oslo2 cohort are available from the Gene Expression Omnibus (GEO) database with accession number GSE84207. For comparison of Oslo2 CpG DNA methylation levels to normal tissue, data from normal breast tissue were available in GEO with accession number GSE60185. The Oslo2 miRNA and mRNA expression data are available from GEO with accession numbers GSE81000 and GSE80999, respectively. For TCGA, the DNA methylation data (level 3) were downloaded from the TCGA Data Portal (https://tcga-data.nci.nih.gov). The miRNA and mRNA expression data (level 3) were downloaded from the UCSC Xena browser [47]. miRNA expression from DCIS samples were available from GEO data set GSE59248 [13]. DNA methylation data from cancer cell lines and immune cells were collected from the GEO with the following accession numbers: GSE94943 (breast cancer cell lines), GSE69270 (leukocytes), GSE68456 (B-cells and monocytes) and GSE79144 (T-cells).

### Competing interests

The authors declare that they have no competing interests.

### Funding

MR Aure was a postdoctoral fellow of the South Eastern Norway Health Authority (grant 2014021 to A-L Børresen-Dale) and a research fellow of the Norwegian Cancer Society (grant 711164 to VN Kristensen). AM and JACM were supported by the Norwegian Research Council (187615); South Eastern Norway Health Authority; and the University of Oslo through the Centre for Molecular Medicine Norway (NCMM); the Norwegian Research Council (288404); and the Norwegian Cancer Society (197884).

### Author’s contributions

Conception and design: MRA, TF, XT, VNK; Development of methodology: MRA, TF, SB, AM, XT; Acquisition of data: MRA, TF, SB, JA, JACM, OSBREAC, ALBD, KKS, AM, XT, VNK; Analysis and interpretation of data: MRA, TF, SB, JA, JACM, AM, XT; Writing of the manuscript: MRA, AM, XT, VNK; Administrative, technical, or material support: OSBREAC, ALBD, KKS, VNK. All authors read and approved the final manuscript.

## Acknowledgments

We would like to thank Daniel Nebdal for excellent technical assistance and advices.

## OSBREAC members

Tone F Bathen (PhD), Norwegian University of Science and Technology, Norway

Elin Borgen (PhD, MD) Oslo University Hospital, Norway

Olav Engebråten (PhD, MD),Oslo University Hospital, Norway

Britt Fritzman (MD), Østfold Hospital, Norway

Øystein Garred (PhD, MD), Oslo University Hospital, Norway

Jürgen Geisler (PhD, MD) Akershus University Hospital, Norway

Gry Aarum Geitvik, Oslo University Hospital, Norway

Solveig Hofvind (PhD),Cancer Registry of Norway, Norway

Rolf Kåresen (PhD, MD),Oslo University Hospital, Norway

Anita Langerød (PhD),Oslo University Hospital, Norway

Ole Christian Lingjærde (PhD), University of Oslo, Norway

Gunhild Mari Mælandsmo (PhD), Oslo University Hospital, Norway

Bjørn Naume (PhD, MD), Oslo University Hospital, Norway

Hege G Russnes (PhD, MD),Oslo University Hospital, Norway

Torill Sauer (PhD, MD),Akershus University Hospital, Norway

Helle Kristine Skjerven (MD), Vestre Viken Hospital Trust, Norway

Ellen Schlichting (PhD, MD), Oslo University Hospital, Norway

Therese Sørlie (PhD),Oslo University Hospital, Norway

## Notes

### Competing Interest Statement

The authors have declared no competing interest.

## References

1. Fleischer T, Frigessi A, Johnson KC, Edvardsen H, Touleimat N, Klajic J, Riis ML, Haakensen VD, Wärnberg F, Naume B, et al: Genome-wide DNA methylation profiles in progression to in situ and invasive carcinoma of the breast with impact on gene transcription and prognosis. Genome Biology 2014, 15:435.

2. Fleischer T, Tekpli X, Mathelier A, Wang S, Nebdal D, Dhakal HP, Sahlberg KK, Schlichting E, Sauer T, Geisler J, et al: DNA methylation at enhancers identifies distinct breast cancer lineages. Nature Communications 2017, 8:1379.

3. Stefansson OA, Moran S, Gomez A, Sayols S, Arribas-Jorba C, Sandoval J, Hilmarsdottir H, Olafsdottir E, Tryggvadottir L, Jonasson JG, et al: A DNA methylation-based definition of biologically distinct breast cancer subtypes. Molecular Oncology 2015, 9:555–568.

4. Achinger-Kawecka J, Valdes-Mora F, Luu P-L, Giles KA, Caldon CE, Qu W, Nair S, Soto S, Locke WJ, Yeo-Teh NS, et al: Epigenetic reprogramming at estrogen-receptor binding sites alters 3D chromatin landscape in endocrine-resistant breast cancer. Nature Communications 2020, 11:320.

5. Rodríguez-Paredes M, Esteller M: Cancer epigenetics reaches mainstream oncology. Nature Medicine 2011, 17:330–339.

6. You Jueng S, Jones Peter A: Cancer Genetics and Epigenetics: Two Sides of the Same Coin? Cancer Cell 2012, 22:9–20.

7. Bartel DP: MicroRNAs: Target Recognition and Regulatory Functions. Cell 2009, 136:215–233.

8. Rupaimoole R, Slack FJ: MicroRNA therapeutics: towards a new era for the management of cancer and other diseases. Nature Reviews Drug Discovery 2017, 16:203.

9. McCall MN, Kim M-S, Adil M, Patil AH, Lu Y, Mitchell CJ, Leal-Rojas P, Xu J, Kumar M, Dawson VL, et al: Toward the human cellular microRNAome. Genome Research 2017, 27:1769–1781.

10. Enerly E, Steinfeld I, Kleivi K, Leivonen S-K, Aure MR, Russnes HG, Rønneberg JA, Johnsen H, Navon R, Rødland E, et al: miRNA-mRNA Integrated Analysis Reveals Roles for miRNAs in Primary Breast Tumors. PLoS ONE 2011, 6:e16915.

11. Blenkiron C, Goldstein L, Thorne N, Spiteri I, Chin S-F, Dunning M, Barbosa-Morais N, Teschendorff A, Green A, Ellis I, et al: MicroRNA expression profiling of human breast cancer identifies new markers of tumor subtype. Genome Biology 2007, 8:R214.

12. The Cancer Genome Atlas Network: Comprehensive molecular portraits of human breast tumours. Nature 2012, 490:61–70.

13. Lesurf R, Aure Miriam R, Mørk Hanne H, Vitelli V, Sauer T, Geisler J, Hofvind S, Borgen E, Børresen-Dale A-L, Engebråten O, et al: Molecular Features of Subtype-Specific Progression from Ductal Carcinoma In Situ to Invasive Breast Cancer. Cell Reports 2016, 16:1166–1179.

14. Haakensen VD, Nygaard V, Greger L, Aure MR, Fromm B, Bukholm IRK, Lüders T, Chin S-F, Git A, Caldas C, et al: Subtype-specific micro-RNA expression signatures in breast cancer progression. International Journal of Cancer 2016, 139:1117–1128.

15. Tahiri A, Leivonen S-K, Lüders T, Steinfeld I, Ragle Aure M, Geisler J, Mäkelä R, Nord S, Riis MLH, Yakhini Z, et al: Deregulation of cancer-related miRNAs is a common event in both benign and malignant human breast tumors. Carcinogenesis 2014, 35:76–85.

16. Dvinge H, Git A, Graf S, Salmon-Divon M, Curtis C, Sottoriva A, Zhao Y, Hirst M, Armisen J, Miska EA, et al: The shaping and functional consequences of the microRNA landscape in breast cancer. Nature 2013, 497:378–382.

17. Aure MR, Vitelli V, Jernström S, Kumar S, Krohn M, Due EU, Haukaas TH, Leivonen S-K, Vollan HKM, Lüders T, et al: Integrative clustering reveals a novel split in the luminal A subtype of breast cancer with impact on outcome. Breast Cancer Research 2017, 19:44.

18. Aure MR, Jernstrom S, Krohn M, Vollan H, Due E, Rodland E, Karesen R, Ram P, Lu Y, Mills G, et al: Integrated analysis reveals microRNA networks coordinately expressed with key proteins in breast cancer. Genome Medicine 2015, 7:21.

19. Fabbri M, Garzon R, Cimmino A, Liu Z, Zanesi N, Callegari E, Liu S, Alder H, Costinean S, Fernandez-Cymering C, et al: MicroRNA-29 family reverts aberrant methylation in lung cancer by targeting DNA methyltransferases 3A and 3B. Proceedings of the National Academy of Sciences 2007, 104:15805–15810.

20. Chen Q, Yin D, Zhang Y, Yu L, Li X-D, Zhou Z-J, Zhou S-L, Gao D-M, Hu J, Jin C, et al: MicroRNA-29a induces loss of 5-hydroxymethylcytosine and promotes metastasis of hepatocellular carcinoma through a TET–SOCS1–MMP9 signaling axis. Cell Death & Disease 2017, 8:e2906–e2906.

21. Aure MR, Leivonen S-K, Fleischer T, Zhu Q, Overgaard J, Alsner J, Tramm T, Louhimo R, Alnæs GIG, Perälä M, et al: Individual and combined effects of DNA methylation and copy number alterations on miRNA expression in breast tumors. Genome Biology 2013, 14:R126.

22. Kuleshov MV, Jones MR, Rouillard AD, Fernandez NF, Duan Q, Wang Z, Koplev S, Jenkins SL, Jagodnik KM, Lachmann A, et al: Enrichr: a comprehensive gene set enrichment analysis web server 2016 update. Nucleic Acids Research 2016, 44:W90–W97.

23. Ju W, Greene CS, Eichinger F, Nair V, Hodgin JB, Bitzer M, Lee Y-s, Zhu Q, Kehata M, Li M, et al: Defining cell-type specificity at the transcriptional level in human disease. Genome Research 2013.

24. Aran D, Hu Z, Butte AJ: xCell: digitally portraying the tissue cellular heterogeneity landscape. Genome Biology 2017, 18:220.

25. Ali HR, Provenzano E, Dawson SJ, Blows FM, Liu B, Shah M, Earl HM, Poole CJ, Hiller L, Dunn JA, et al: Association between CD8+ T-cell infiltration and breast cancer survival in 12 439 patients. Annals of Oncology 2014, 25:1536–1543.

26. Tekpli X, Lien T, Røssevold AH, Nebdal D, Borgen E, Ohnstad HO, Kyte JA, Vallon-Christersson J, Fongaard M, Due EU, et al: An independent poor-prognosis subtype of breast cancer defined by a distinct tumor immune microenvironment. Nature Communications 2019, 10:5499.

27. Walker C, Mojares E, Del Río Hernández A: Role of Extracellular Matrix in Development and Cancer Progression. International Journal of Molecular Sciences 2018, 19:3028.

28. Gheorghe M, Sandve GK, Khan A, Chèneby J, Ballester B, Mathelier A: A map of direct TF–DNA interactions in the human genome. Nucleic Acids Research 2018, 47:e21–e21.

29. Xi Y, Shi J, Li W, Tanaka K, Allton KL, Richardson D, Li J, Franco HL, Nagari A, Malladi VS, et al: Histone modification profiling in breast cancer cell lines highlights commonalities and differences among subtypes. BMC Genomics 2018, 19:150.

30. Wu VT, Kiriazov B, Koch KE, Gu VW, Beck AC, Borcherding N, Li T, Addo P, Wehrspan ZJ, Zhang W, et al: A TFAP2C Gene Signature Is Predictive of Outcome in HER2-Positive Breast Cancer. Molecular Cancer Research 2020, 18:46–56.

31. Suzuki HI, Young RA, Sharp PA: Super-Enhancer-Mediated RNA Processing Revealed by Integrative MicroRNA Network Analysis. Cell 2017, 168:1000-1014.e1015.

32. Li G, Ruan X, Auerbach Raymond K, Sandhu Kuljeet S, Zheng M, Wang P, Poh Huay M, Goh Y, Lim J, Zhang J, et al: Extensive Promoter-Centered Chromatin Interactions Provide a Topological Basis for Transcription Regulation. Cell 2012, 148:84–98.

33. Li D, Hsu S, Purushotham D, Sears RL, Wang T: WashU Epigenome Browser update 2019. Nucleic Acids Research 2019, 47:W158–W165.

34. Friedman RC, Farh KK-H, Burge CB, Bartel DP: Most mammalian mRNAs are conserved targets of microRNAs. Genome Research 2009, 19:92–105.

35. Johansson HJ, Socciarelli F, Vacanti NM, Haugen MH, Zhu Y, Siavelis I, Fernandez-Woodbridge A, Aure MR, Sennblad B, Vesterlund M, et al: Breast cancer quantitative proteome and proteogenomic landscape. Nature Communications 2019, 10:1600.

36. Pietras K, Östman A: Hallmarks of cancer: Interactions with the tumor stroma. Experimental Cell Research 2010, 316:1324–1331.

37. Nagpal V, Rai R, Place AT, Murphy SB, Verma SK, Ghosh AK, D. V: MiR-125b Is Critical for Fibroblast-to-Myofibroblast Transition and Cardiac Fibrosis. Circulation 2016, 133:291–301.

38. Hu Y, Zhu Q, Tang L: MiR-99a Antitumor Activity in Human Breast Cancer Cells through Targeting of mTOR Expression. PLOS ONE 2014, 9:e92099.

39. Feliciano A, Castellvi J, Artero-Castro A, Leal JA, Romagosa C, Hernández-Losa J, Peg V, Fabra A, Vidal F, Kondoh H, et al: miR-125b Acts as a Tumor Suppressor in Breast Tumorigenesis via Its Novel Direct Targets ENPEP, CK2-α, CCNJ, and MEGF9. PLOS ONE 2013, 8:e76247.

40. Khan S, Brougham CL, Ryan J, Sahrudin A, O’Neill G, Wall D, Curran C, Newell J, Kerin MJ, Dwyer RM: miR-379 Regulates Cyclin B1 Expression and Is Decreased in Breast Cancer. PLOS ONE 2013, 8:e68753.

41. Lee Y, Ahn C, Han J, Choi H, Kim J, Yim J, Lee J, Provost P, Radmark O, Kim S, Kim VN: The nuclear RNase III Drosha initiates microRNA processing. Nature 2003, 425:415–419.

42. Li Y, Zhang Y, Li S, Lu J, Chen J, Wang Y, Li Y, Xu J, Li X: Genome-wide DNA methylome analysis reveals epigenetically dysregulated non-coding RNAs in human breast cancer. Scientific Reports 2015, 5:8790.

43. Chou J, Lin JH, Brenot A, Kim J-w, Provot S, Werb Z: GATA3 suppresses metastasis and modulates the tumour microenvironment by regulating microRNA-29b expression. Nature Cell Biology 2013, 15:201–213.

44. Li W, Yi J, Zheng X, Liu S, Fu W, Ren L, Li L, Hoon DSB, Wang J, Du G: miR-29c plays a suppressive role in breast cancer by targeting the TIMP3/STAT1/FOXO1 pathway. Clinical Epigenetics 2018, 10:64.

45. Fleischer T, Klajic J, Aure MR, Louhimo R, Pladsen AV, Ottestad L, Touleimat N, Laakso M, Halvorsen AR, Alnæs GIG, et al: DNA methylation signature (SAM40) identifies subgroups of the Luminal A breast cancer samples with distinct survival. Oncotarget 2017, 8:1074–1082.

46. Touleimat N, Tost J: Complete pipeline for Infinium® Human Methylation 450K BeadChip data processing using subset quantile normalization for accurate DNA methylation estimation. Epigenomics 2012, 4:325–341.

47. Goldman M, Craft B, Hastie M, Repečka K, Kamath A, McDade F, Rogers D, Brooks AN, Zhu J, Haussler D: The UCSC Xena platform for public and private cancer genomics data visualization and interpretation. bioRxiv 2019:326470.

48. The R Development Core Team: R: A language and environment for statistical computing. R Foundation for Statistical Computing, Vienna, Austria.; 2011.

49. Kolde R: pheatmap: Pretty Heatmaps. R package version 1.0.12. https://CRAN.R-project.org/package=pheatmap. 2019.

50. Benjamini Y, Hochberg Y: Controlling the False Discovery Rate - a Practical and Powerful Approach to Multiple Testing. Journal of the Royal Statistical Society Series B-Methodological 1995, 57:289 – 300.

51. Quinlan AR, Hall IM: BEDTools: a flexible suite of utilities for comparing genomic features. Bioinformatics 2010, 26:841–842.

52. Fornes O, Castro-Mondragon JA, Khan A, van der Lee R, Zhang X, Richmond PA, Modi BP, Correard S, Gheorghe M, Baranašić D, et al: JASPAR 2020: update of the open–access database of transcription factor binding profiles. Nucleic Acids Research 2020, 48:D87–D92.

53. Sheffield NC, Bock C: LOLA: enrichment analysis for genomic region sets and regulatory elements in R and Bioconductor. Bioinformatics 2016, 32:587–589.

54. Chèneby J, Gheorghe M, Artufel M, Mathelier A, Ballester B: ReMap 2018: an updated atlas of regulatory regions from an integrative analysis of DNA-binding ChIP-seq experiments. Nucleic Acids Research 2017, 46:D267–D275.

55. Lawrence M, Huber W, Pagès H, Aboyoun P, Carlson M, Gentleman R, Morgan MT, Carey VJ: Software for Computing and Annotating Genomic Ranges. PLOS Computational Biology 2013, 9:e1003118.

