## Additional files 1-2, 5-14 for "Crosstalk between microRNA expression and DNA methylation drive the hormone-dependent phenotype of breast cancer"

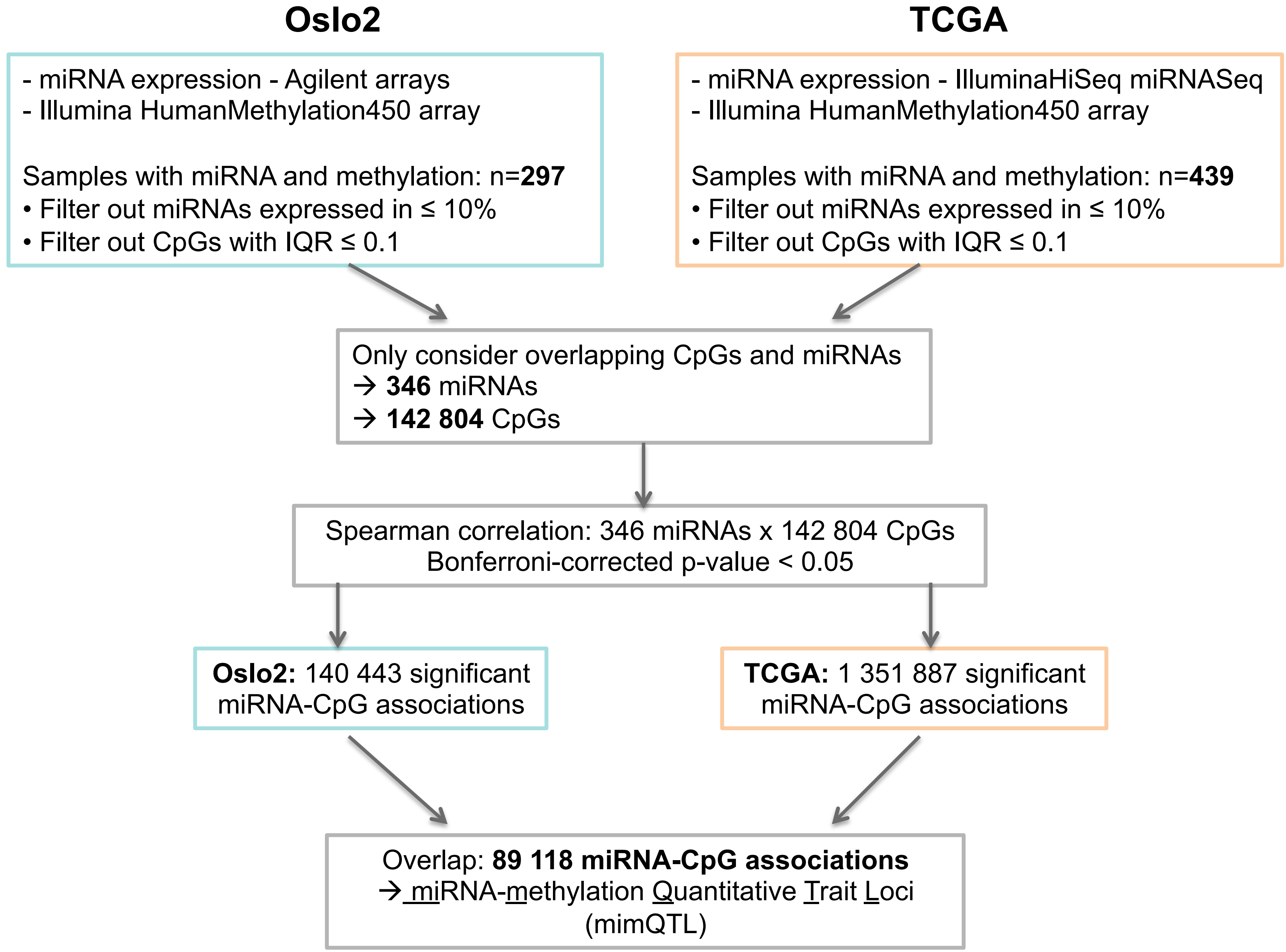


**Additional file 1.** Flowchart describing the different steps of the analysis leading to the identification of 89 118 miRNA-methylation Quantitative Trait Loci (mimQTLs).

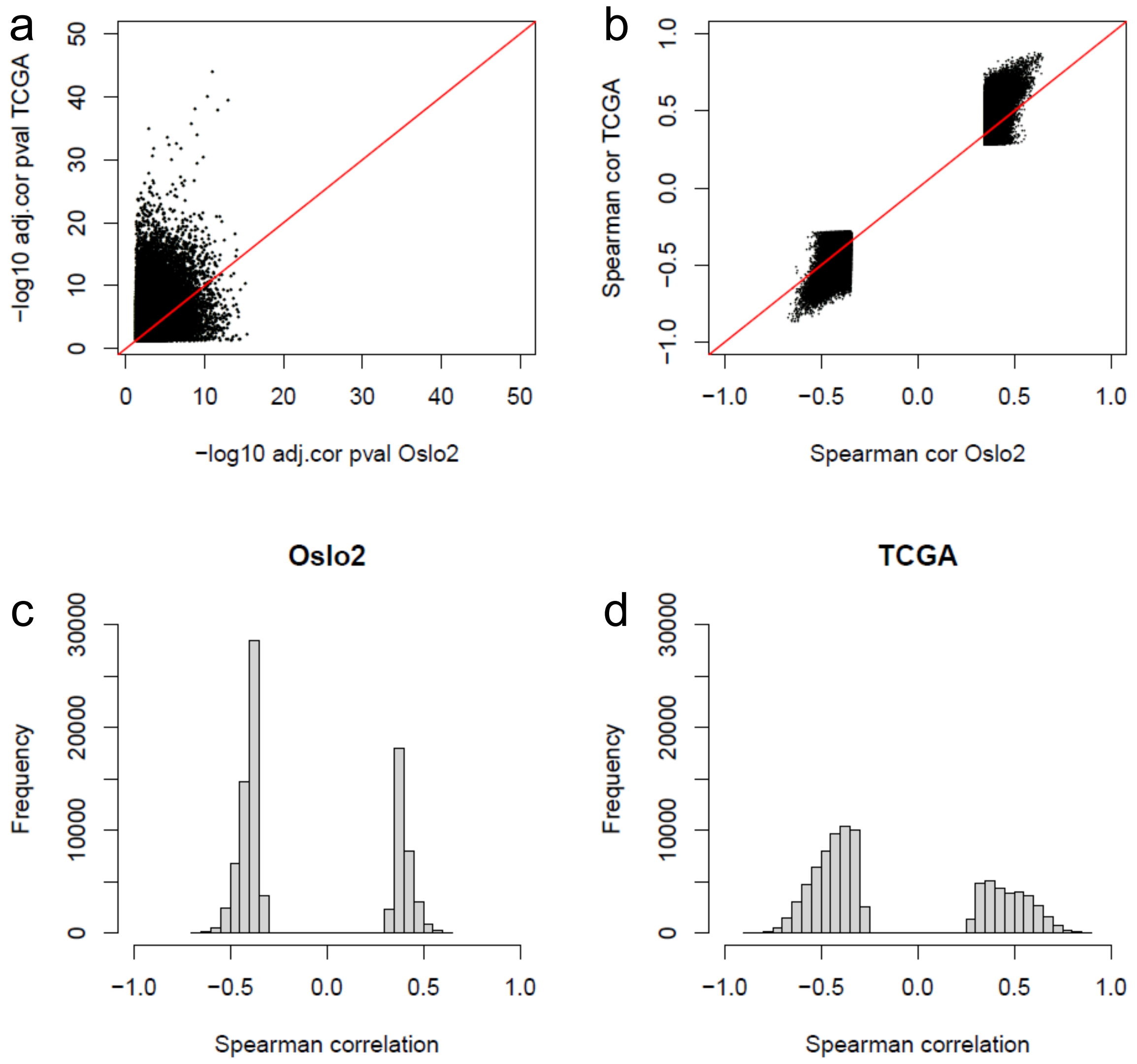


**Additional file 2.** Overview of the 89 118 miRNA-CpG associations found significant in both cohorts. The scatterplots show **a)** the –log10(Bonferroni adjusted Spearman correlation p-values) of Oslo2 vs. TCGA; **b)** Spearman correlation coefficients in Oslo2 vs. TCGA. The histograms show the distribution of the correlation coefficients (Spearman's rho) of all significant miRNA-CpG correlations in **c)** Oslo2 and **d)** TCGA cohorts.

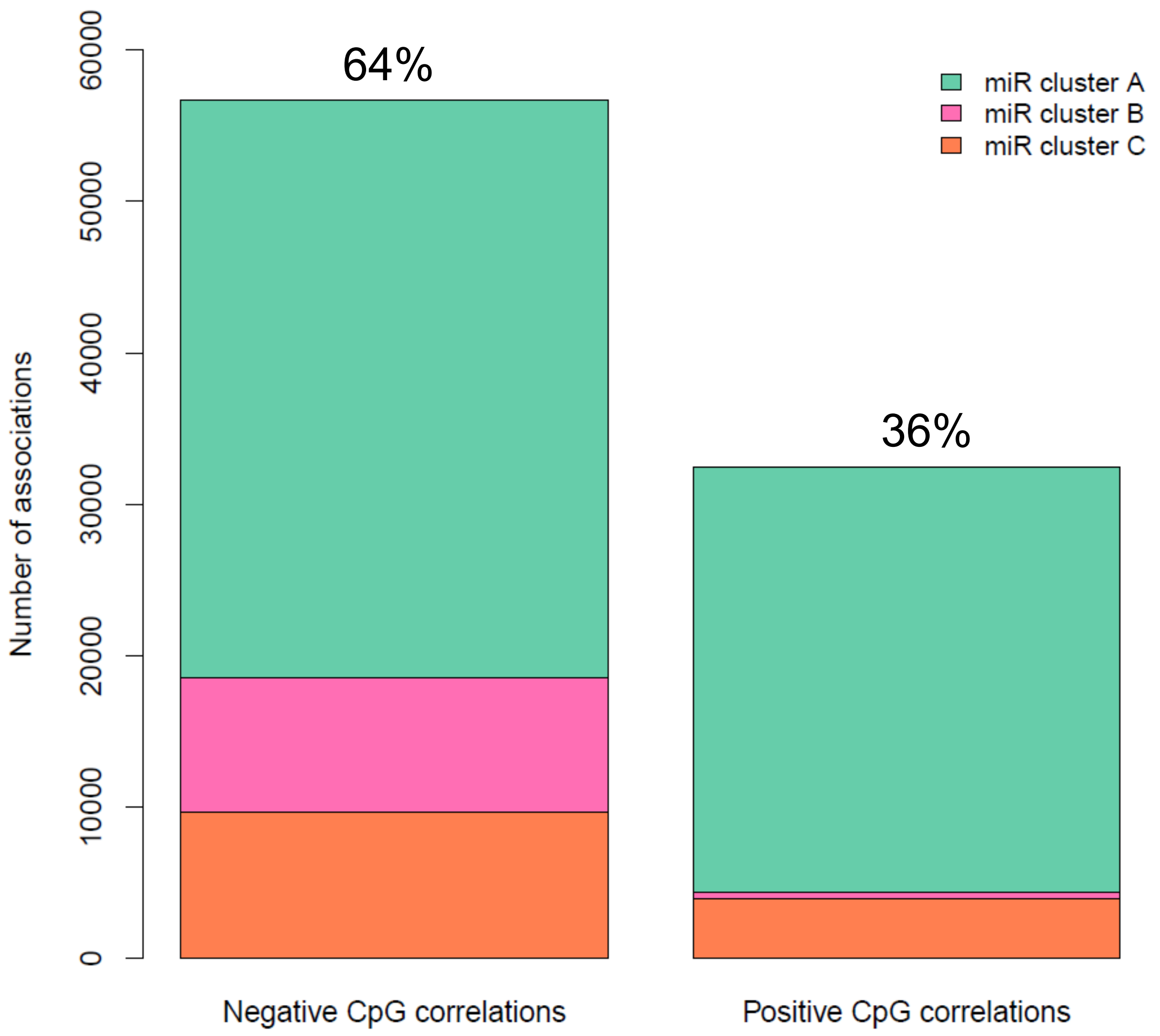


**Additional file 5.** Barplots showing the number of negative and positive CpG correlations for the three different miRNA clusters.

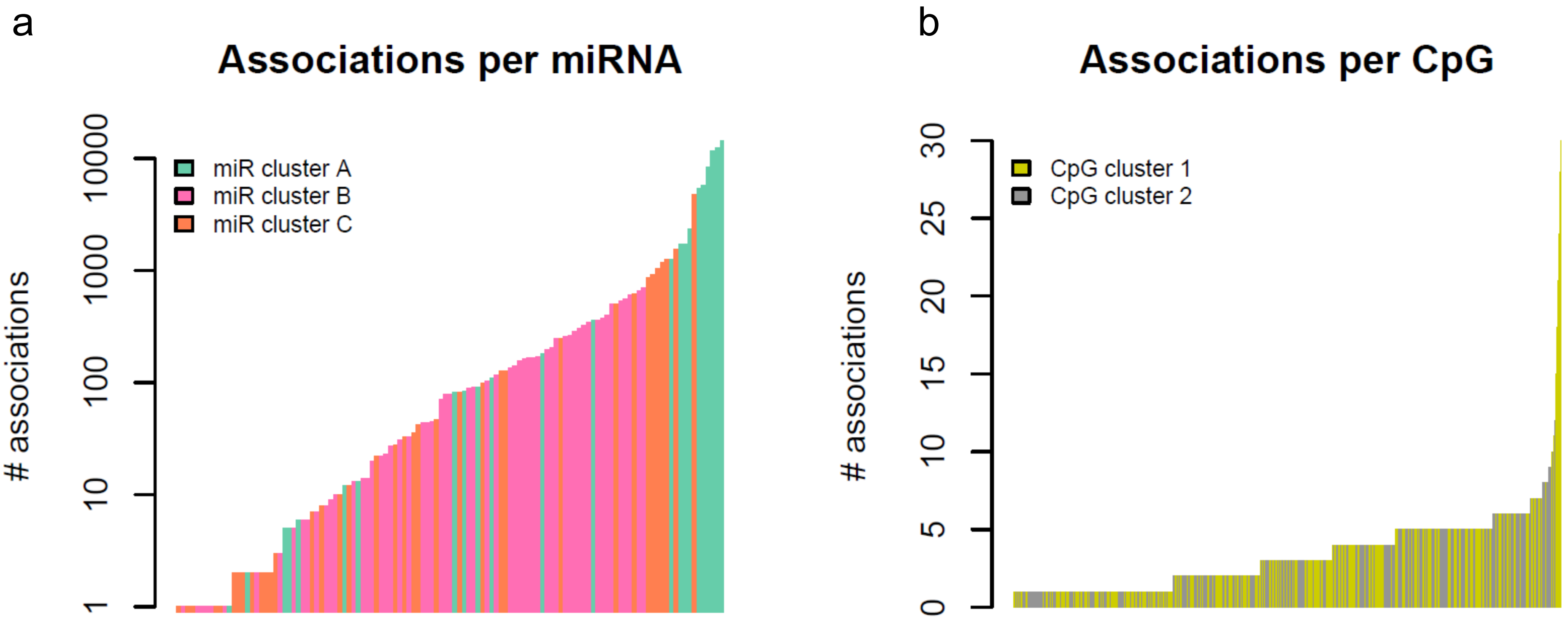


**Additional file 6.** Barplots showing number of associations per miRNA or CpG. **a)** Number of CpG associations per miRNA (n=119). Note that the y-axis is on log scale. **b)** Number of miRNA associations per CpG (n=26746).

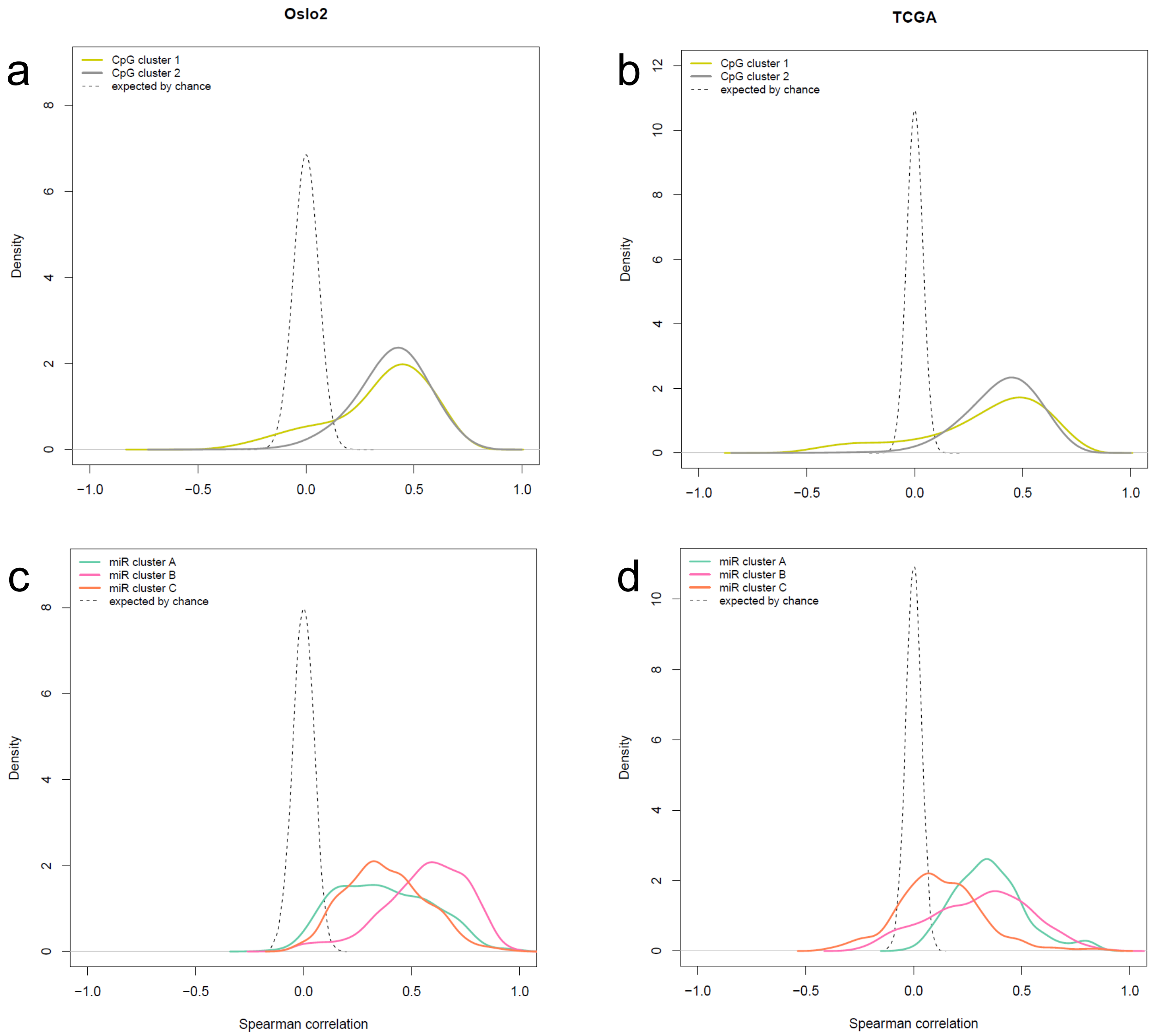


**Additional file 7.** Density plots showing the degree of miRNA co-expression or CpG co-methylation between cluster members (see Figure 1) calculated by Spearman correlation. **a)** Correlation between CpG cluster members in the Oslo2 data. **b)** Correlation between CpG cluster members in the TCGA data. **c)** Correlation between miRNA cluster members in the Oslo2 data. **d)** Correlation between miRNA cluster members in the TCGA data. The dotted lines represent density plots of corresponding correlations expected by chance, i.e. correlations observed after randomly permuting the same data before performing correlation analyses.

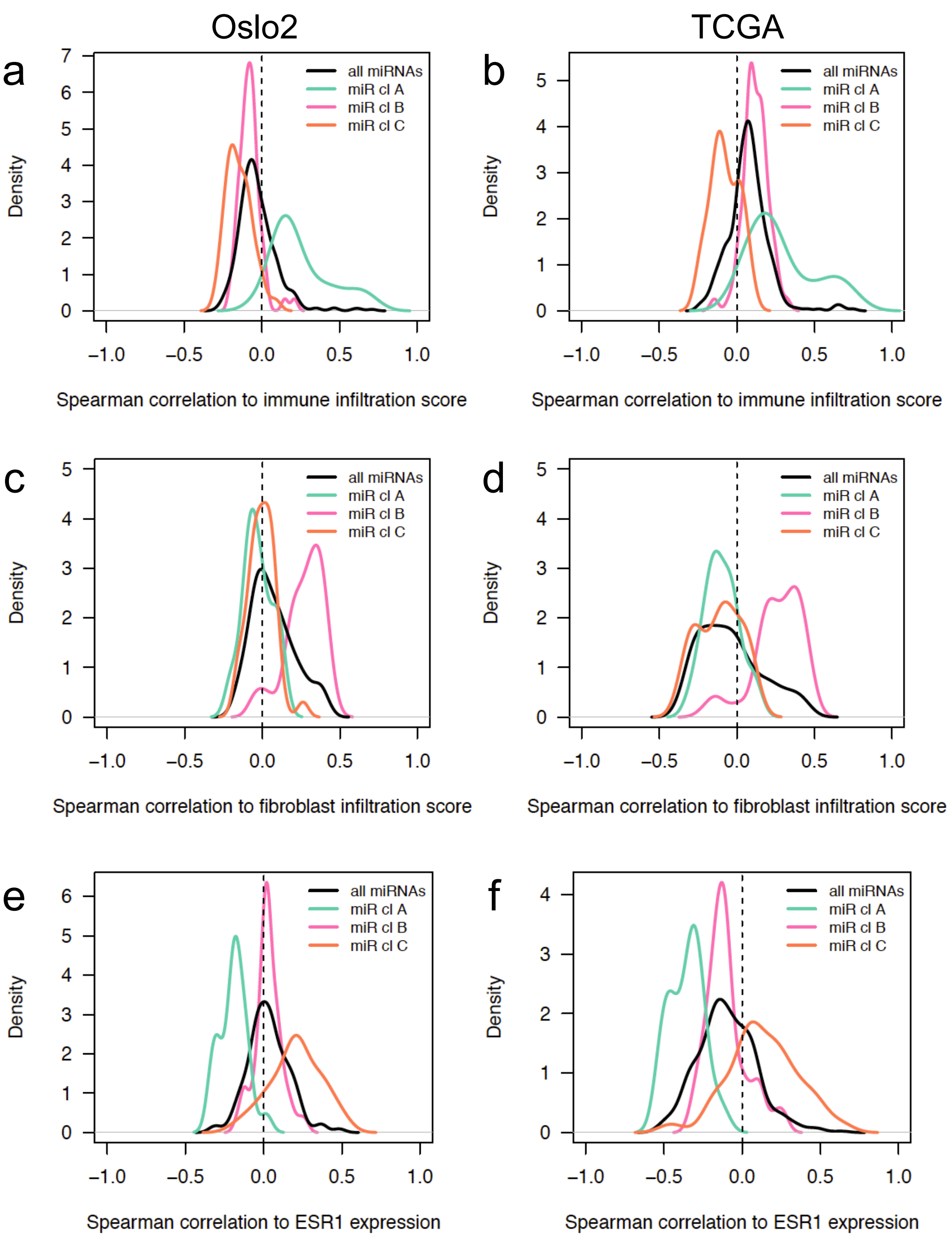


**Additional file 8.** Density plots showing the distribution of Spearman correlation coefficients between miRNA expression and selected variables for members of each of the miRNA clusters. **a, b)** miRNA expression-immune infiltration score [23] correlations for the Oslo2 **(a)** and TCGA cohorts **(b)**. **c, d)** miRNA expression-fibroblast infiltration score [24] correlations for the Oslo2 **(c)** and TCGA cohorts **(d)**. **e, f)** miRNA expression-ESR1 mRNA expression correlations for the Oslo2 **(e)** and TCGA cohorts **(f).**

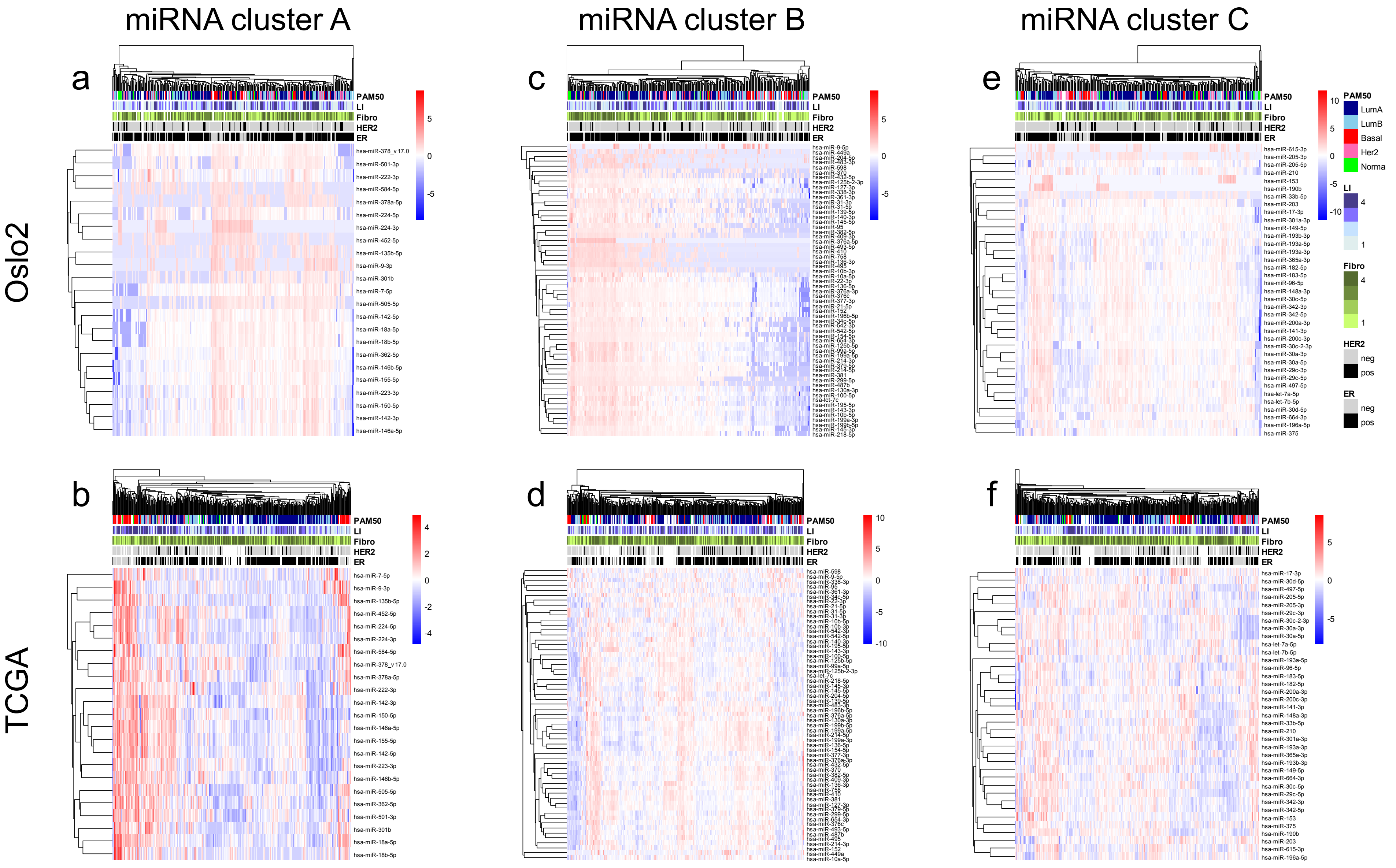


**Additional file 9.** Heatmaps showing hierarchical clustering of miRNA expression levels (rows) from tumors (columns) of the Oslo2 (top) and TCGA (bottom) cohort. Clustering was performed using Euclidean distance and average linkage. Tumors are annotated with PAM50 molecular subtypes (Luminal A (LumA), Luminal B (LumB), Basal-like (Basal), HER2-enriched (Her2), Normal-like (Normal); Lymphocyte infiltration (LI) group where tumors were divided into quartiles: 1 (low) – 4 (high); Fibroblast infiltration group (Fibro) where tumors were divided into quartiles: 1 (low) – 4 (high); Human epidermal growth factor receptor 2 (HER2) status; Estrogen receptor (ER) status. **a, b)** Clustering of miRNA cluster A expression (n=23); **c, d)** Clustering of miRNA cluster B expression (n=59); **e, f)** Clustering of miRNA cluster C expression (n=37).

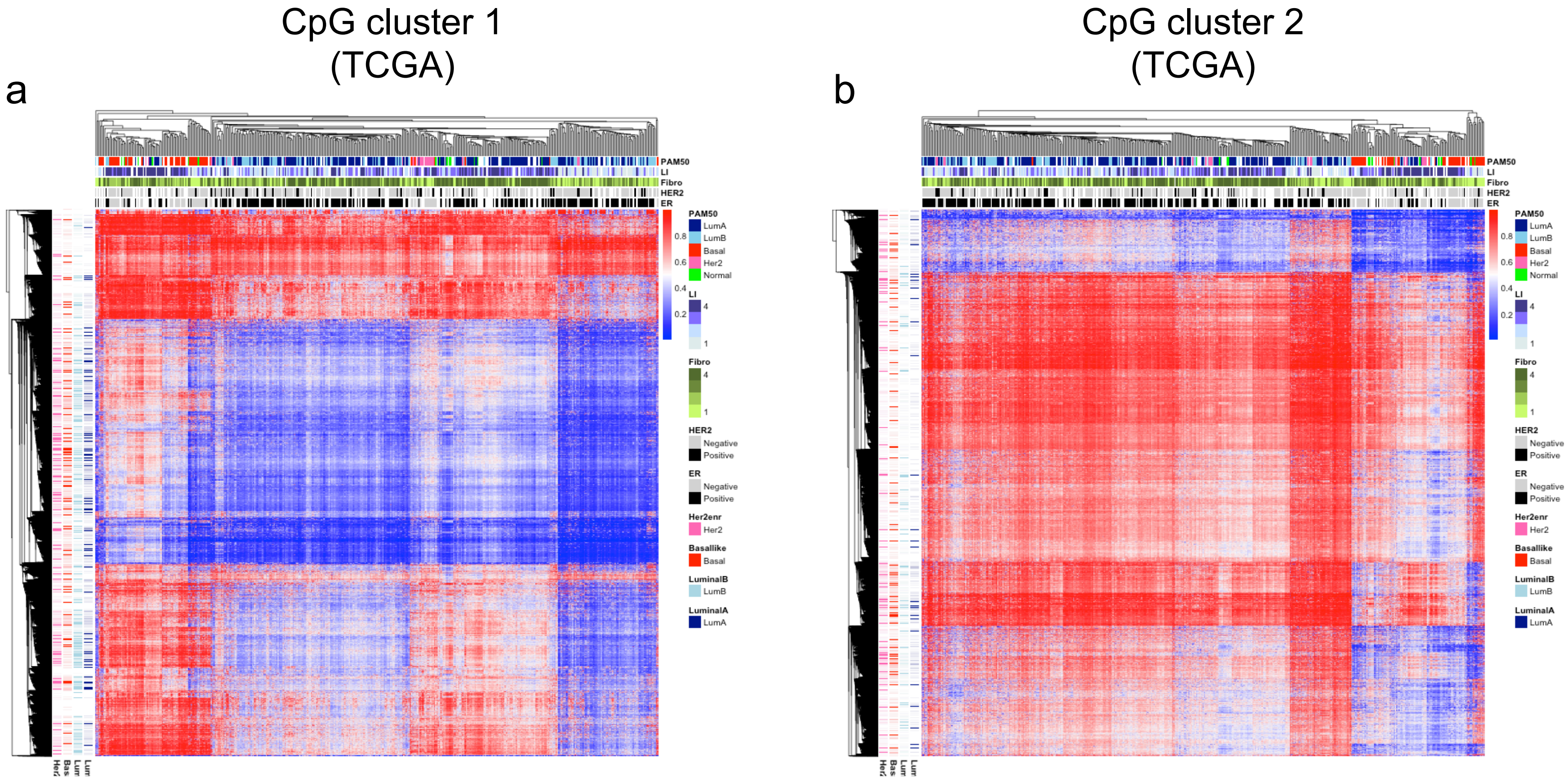


**Additional file 10.** Heatmaps showing hierarchical clustering of methylation levels of CpG cluster 1 (**a**; n = 14040) and CpG cluster 2 (**b;** n=12706) in the TCGA cohort (CpGs in rows and tumors in columns). Clustering was performed using Euclidean distance and average linkage. Tumors are annotated with the following clinical/ molecular classifications: PAM50 molecular subtypes (Luminal A (LumA), Luminal B (LumB), Basal-like (Basal), HER2-enriched (Her2), Normal-like (Normal); Lymphocyte infiltration (LI) group where tumors were divided into quartiles: 1 (low) – 4 (high); Human epidermal growth factor receptor 2 (HER2) status; Estrogen receptor (ER) status. The CpGs are annotated according to overlap with regions annotated as “active intergenic enhancer” from ChromHMM of subtype-specific cell lines [29].

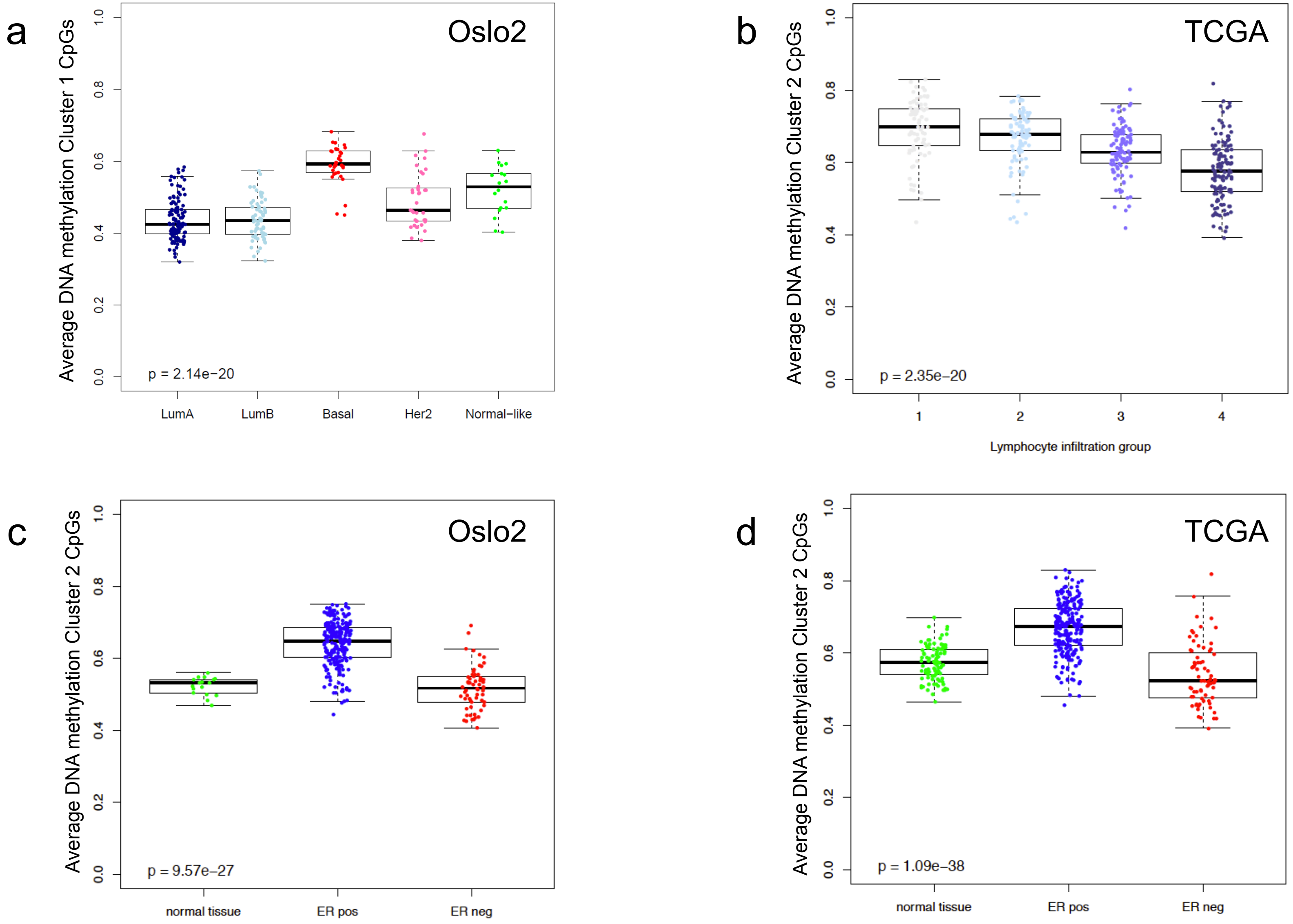


**Additional file 11. a)** Boxplot showing average DNA methylation of CpGs from cluster 1 in PAM50 subtypes of the Oslo2 cohort (Luminal A (LumA), Luminal B (LumB), Basal-like (Basal), HER2-enriched (Her2)). **b)** Boxplot showing average DNA methylation of CpGs from cluster 2 in the TCGA cohort when tumors were separated into quartile lymphocyte infiltration groups from low (1) to high (4) infiltration. **c)** Boxplot showing average DNA methylation of CpGs from cluster 2 in normal breast tissue (reduction mammoplasty, n=17) or estrogen receptor (ER) positive (pos) or negative (neg) tumors of the Oslo2 cohort. **d)** Boxplot showing average DNA methylation of CpGs from cluster 2 in normal breast tissue (normal adjacent breast tissue, n=97) or ER positive or negative tumors of the TCGA cohort. P-values resulting from Kruskal-Wallis tests indicated.

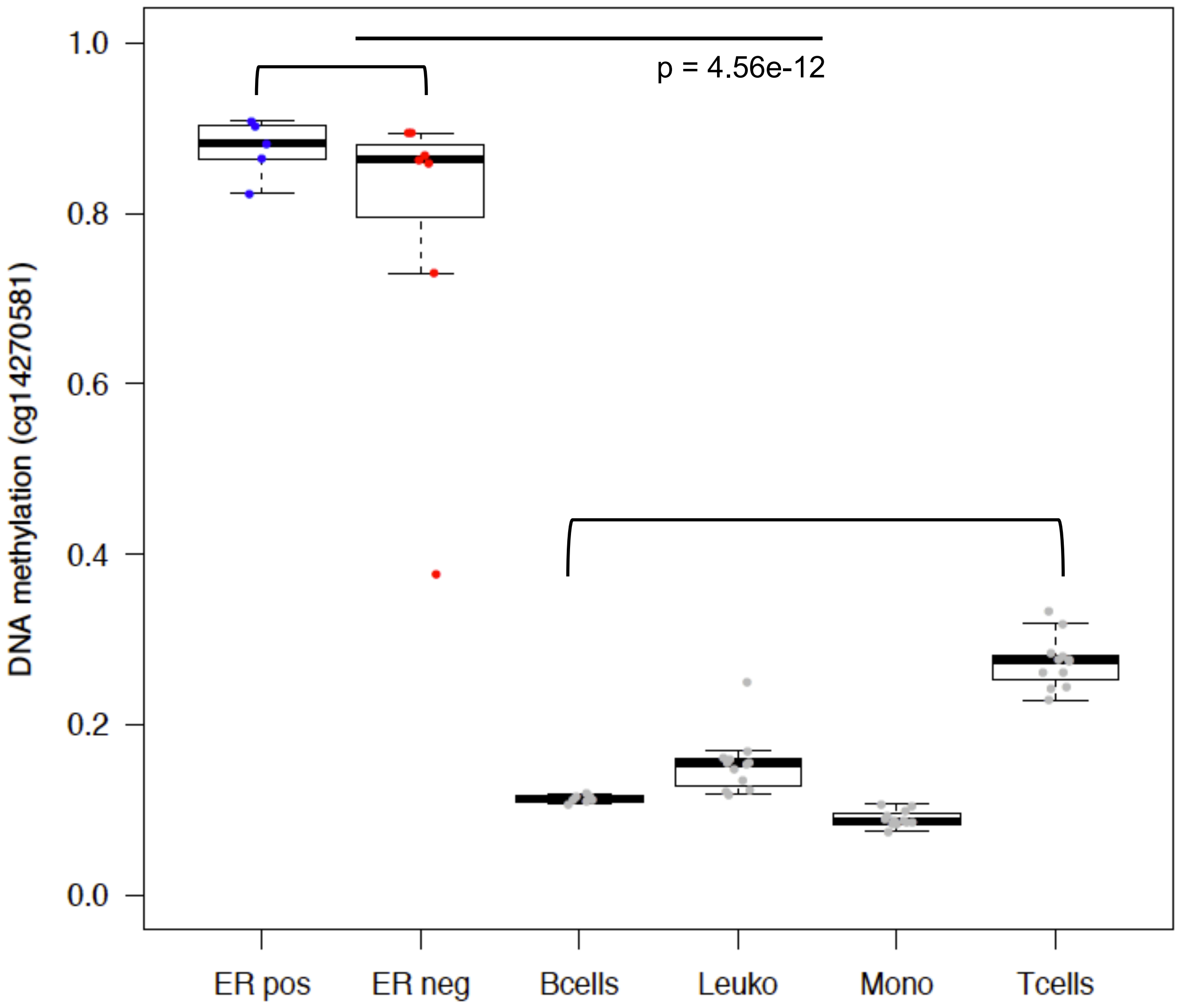


**Additional file 12.** Boxplot showing DNA methylation of the hub CpG of miRNA cluster A (cg14270581; y-axis)) in ER positive (pos) and negative (neg) breast cancer cell lines and from different immune cell types (x-axis); B-cells, leukocytes (leuko), monocytes (mono) and T-cells. P-value resulting from Wilcoxon rank-sum test between cancer cell lines vs. immune cells is indicated.

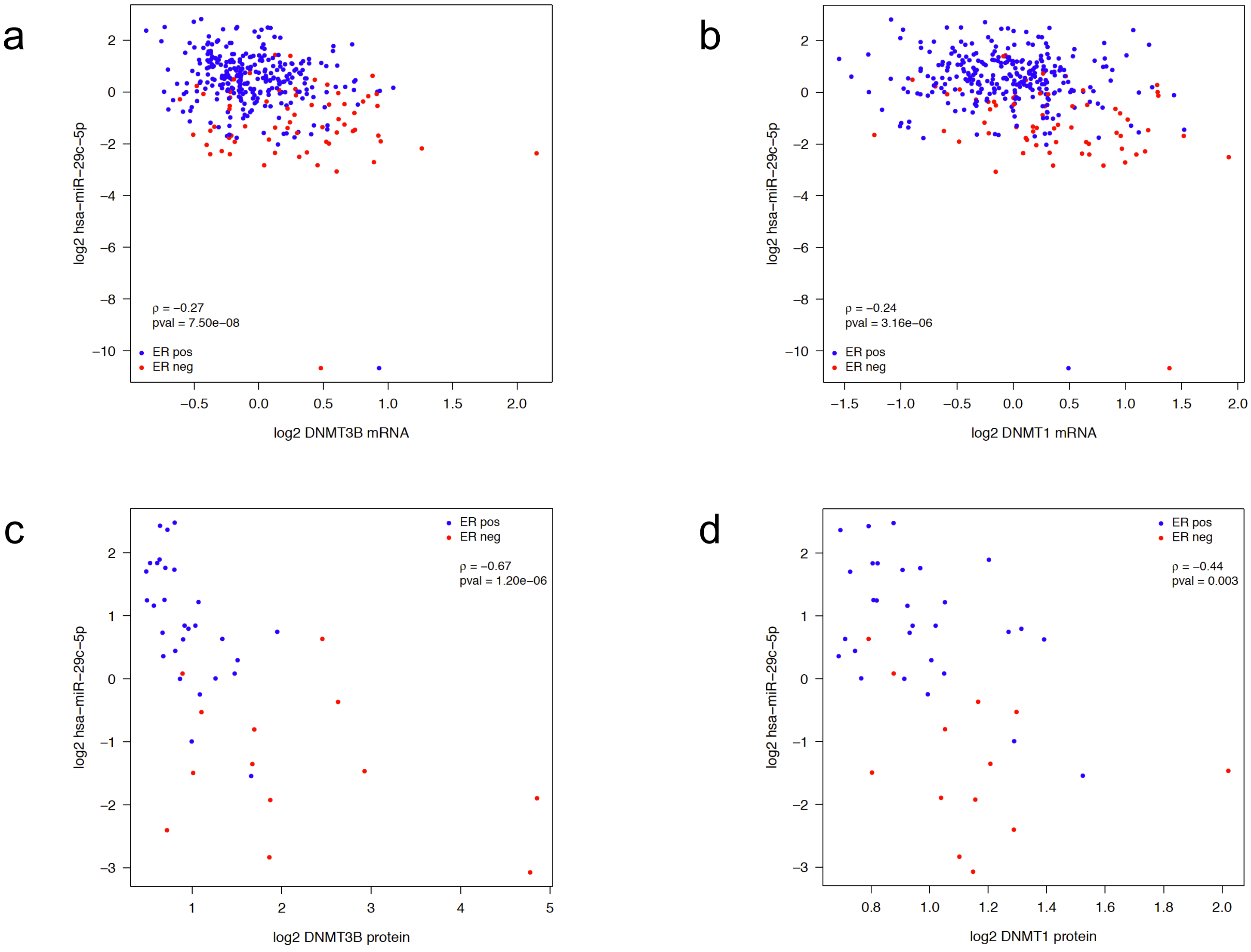


**Additional file 13. a)** DNMT3B mRNA expression (x-axis) vs. hsa-miR-29c-5p expression (y-axis) measured in 377 samples of the Oslo2 cohort. **b)** DNMT1 mRNA expression (x-axis) vs. hsa-miR-29c-5p expression (y-axis) measured in 377 samples of the Oslo2 cohort. **c)** DNMT3B protein expression (x-axis) vs. hsa-miR-29c-5p expression (y-axis) measured in 45 samples of the Oslo2 cohort. **d)** DNMT1 protein expression (x-axis) vs. hsa-miR-29c-5p expression (y-axis) measured in 45 samples of the Oslo2 cohort. Each dot represent a tumor color-coded according estrogen receptor (ER) status; blue: ER positive (pos), red: ER negative (neg). Spearman correlation indicated.

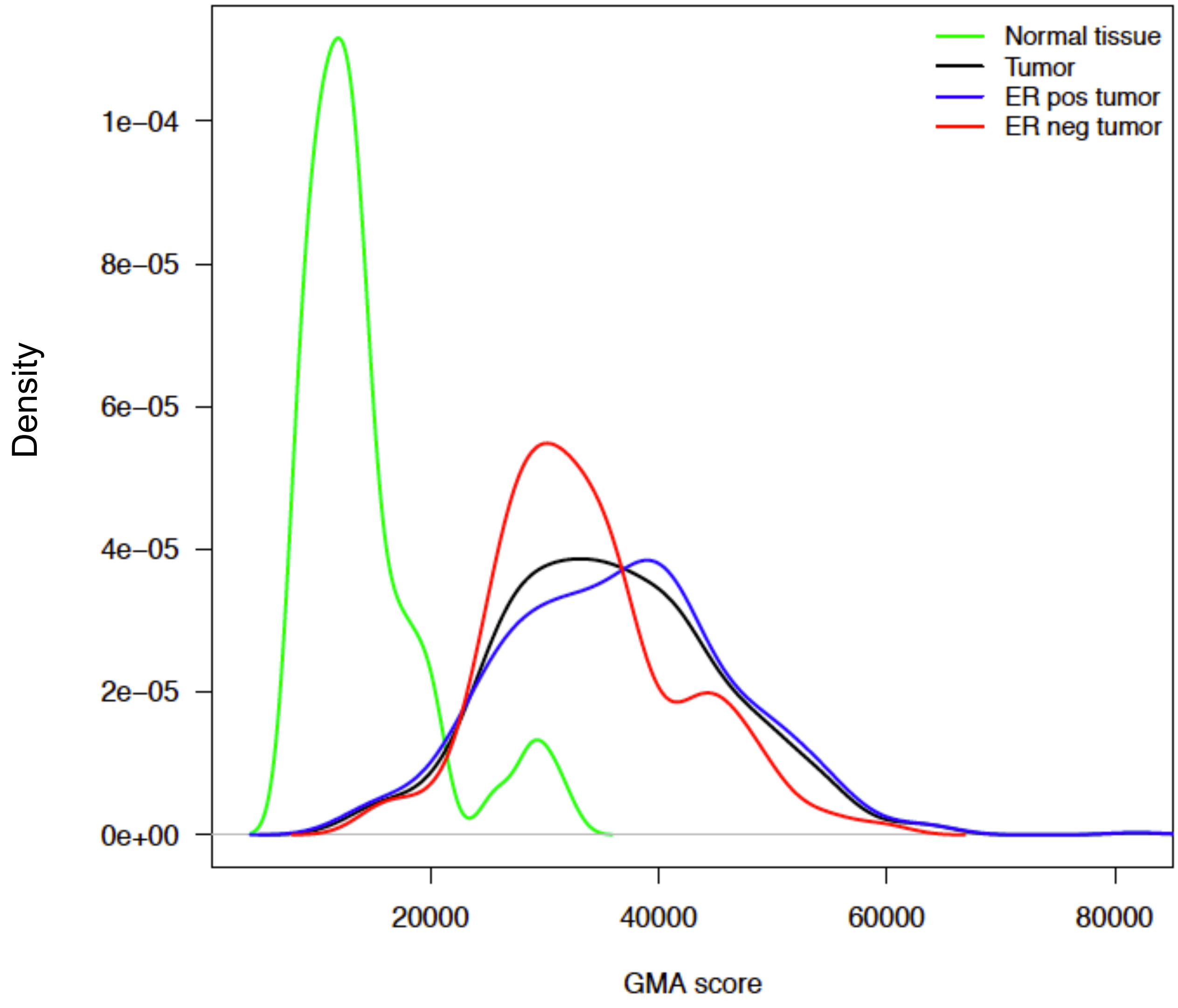


**Additional file 14.** Density plot showing the distribution of the Global Methylation Alteration (GMA) score in normal adjacent breast tissue (green), tumors (black) and tumors separated into estrogen receptor (ER) positive (pos) and negative (neg). Data from TCGA.
